# Molecular architecture of nucleosome remodeling and deacetylase sub-complexes by integrative structure determination

**DOI:** 10.1101/2021.11.25.469965

**Authors:** Shreyas Arvindekar, Matthew J. Jackman, Jason K.K. Low, Michael J. Landsberg, Joel P. Mackay, Shruthi Viswanath

## Abstract

The Nucleosome Remodeling and Deacetylase (NuRD) complex is a chromatin-modifying assembly that regulates gene expression and DNA damage repair. Despite its importance, limited structural information describing the complete NuRD complex is available and a detailed understanding of its mechanism is therefore lacking. Drawing on information from SEC-MALLS, DIA-MS, XLMS, negative-stain EM, X-ray crystallography, NMR spectroscopy, secondary structure predictions and homology models, we applied Bayesian integrative structure determination to investigate the molecular architecture of three NuRD sub-complexes: MTA1-HDAC1-RBBP4 (MHR), MTA1^N^-HDAC1-MBD3^GATAD2CC^ (MHM), and MTA1-HDAC1-RBBP4-MBD3-GATAD2A (NuDe). The integrative structures were corroborated by examining independent crosslinks, cryo-EM maps, biochemical assays, known cancer-associated mutations, and structure predictions from AlphaFold. The robustness of the models was assessed by jack-knifing. Localization of the full-length MBD3, which connects the deacetylase and chromatin remodeling modules in NuRD, has not previously been possible; our models indicate two different locations for MBD3, suggesting a mechanism by which MBD3 in the presence of GATAD2A asymmetrically bridges the two modules in NuRD. Further, our models uncovered three previously unrecognized subunit interfaces in NuDe: HDAC1^C^-MTA1^BAH^, MTA1^BAH^-MBD3^MBD^, and HDAC1^60-100^-MBD3^MBD^. Our approach also allowed us to localize regions of unknown structure, such as HDAC1^C^ and MBD3^IDR^, thereby resulting in the most complete and robustly cross-validated structural characterization of these NuRD sub-complexes so far.

## Introduction

The Nucleosome Remodeling and Deacetylase (NuRD) complex is a multi-protein chromatin-modifying assembly that is expressed in most metazoan tissues and conserved across animals (Basta and Rauchman, 2017; Denslow and Wade, 2007). It regulates gene expression and DNA damage repair by modulating nucleosome accessibility in enhancers and promoters for transcription factors and RNA polymerases (Bornelöv et al., 2018; Li and Kumar, 2010; Reynolds et al., 2013; Smeenk et al., 2010; Yoshida et al., 2008). Subunits of NuRD are implicated in human cancers and various congenital defects (Basta and Rauchman, 2015; Toh and Nicolson, 2009). Considerable diversity is observed in subunit isoforms and NuRD-associated factors across tissues (Burgold et al., 2019; Hoffmann and Spengler, 2019).

NuRD comprises two catalytic modules – a histone deacetylase module and ATP-dependent chromatin-remodeling module (Denslow and Wade, 2007; Low et al., 2020). The deacetylase module contains metastasis-associated proteins (MTA1/2/3) that form a dimeric scaffold for the histone deacetylases (HDAC1/2). It also contains the proteins RBBP4/7, which mediate interactions of NuRD with histone tails and transcription factors (Hong et al., 2005; Lejon et al., 2011). The chromatin-remodeling module contains methyl-CpG DNA-binding proteins (MBD2/3) that recruit NuRD to methylated and/or hemi-methylated DNA, GATA-type zinc-finger proteins (GATAD2A/B), and an ATP-dependent DNA translocase (CHD3/4/5) (Burgold et al., 2019; Low et al., 2020). Note that each of these subunits exists as two or more paralogues that can be incorporated into NuRD; for simplicity, we generally refer to only one paralogue of each subunit herein.

Some structural information is available for the complex. The stoichiometry of the complex, based on a consensus of several recent structural, biochemical and quantitative mass spectrometry studies (Zhang 2016, Millard 2013, Low 2020) appears to be 2:2:4:1:1:1 (MTA:HDAC:RBBP:MBD:GATAD2:CHD). These data also suggest that NuRD can be thought of as two ‘modules’ with distinct enzymatic activities: a deacetylase sub-complex comprising MTA1, HDAC2 and RBBP4 (MHR) and a chromatin remodelling sub-complex comprising CHD4, GATAD2A and MBD3 (MGC). It is notable that MHR also appears to exist in the cell as a distinct assembly, as shown in work that demonstrates an interaction between this sub-complex and the chromatin regulator PWWP2A (Link et al., 2018; Pünzeler et al., 2017).

Experimental structures of parts of the NuRD complex, including the 2:2 MTA1^ELM2/SANT^-HDAC1 dimer, RBBP4 bound to the C-terminal half of MTA1, the MBD domain of MBD3, the coiled-coil dimer of MBD2 and GATAD2A, and CHD4 bound to a nucleosome have been determined by X-ray crystallography, cryo-electron microscopy and NMR spectroscopy (Alqarni et al., 2014; Cramer et al., 2014; Farnung et al., 2020; Gnanapragasam et al., 2011; Millard et al., 2016, 2013). Structures of several other NuRD sub-complexes have also been characterized at various resolutions by negative-stain electron microscopy, including MHR, as well as two simplified assemblies that are not known to have independent biological functions: (i) a 2:2:2 MTA1^N^-HDAC1-MBD3^GATAD2CC^ (MHM) complex, and (ii) a 2:2:4:1:1 MTA1-HDAC1-RBBP4-MBD3-GATAD2A (NuDe complex) (Low et al., 2020; Millard et al., 2020, 2016).

Pairwise interactions between domains and subunits within the MHR, MHM, NuDe, and the endogenous NuRD complexes have also been characterized by chemical crosslinking and mass spectrometry (XLMS) (Kloet et al., 2015; Le Guezennec et al., 2006; Low et al., 2020; Millard et al., 2016; Smits et al., 2013; Spruijt et al., 2021, 2010). A model of the MHM sub-complex, based on crosslink-driven rigid-body docking of known atomic structures together with manual placement of a pair of MTA1-(RBBP4)_2_ structures, has also been reported (Low et al., 2020). While this represents the most complete model of NuRD architecture, it was created manually and accounts for only 30% of residues in the NuRD complex. In fact, only 50% of residues in NuRD have known or readily modeled atomic structures, and the structures of proteins such as MBD3, CHD4, and GATAD2A are largely uncharacterized. While more recent artificial intelligence-based methods such as AlphaFold hold more promise for modelling these uncharacterised regions, significant limitations remain for the modelling of multi-protein complexes, including size limitations and issues with the modelling of disordered or irregularly structured regions in protein assemblies (Jumper et al., 2021). These issues, combined with variability in the paralogue composition and significant structural dynamics, suggest that the full structure of the NuRD complex is likely to remain a challenge in the near future.

A striking feature of the NuRD complex is its asymmetric stoichiometry. The MHR complex displays pseudo two-fold symmetry and yet engages with a 1:1:1 MGC complex. It is known that MBD3 binds to the MHR complex and that the 2:2:4:1:1 MTA1-HDAC1-RBBP4-MBD3^-^ GATAD2A (NuDe complex) is also asymmetric (Low et al., 2020). In contrast, the MHM sub-complex, containing only HDAC1, MBD3 and MTA1, has a 2:2:2 stoichiometry, suggesting that it is GATAD2A that introduces the asymmetry. However, the mechanism by which this asymmetry is introduced in NuDe/NuRD is not known. Also, the structure of full-length MBD3, which contains a significant intrinsically disordered region (IDR; MBD3^71-213^) critical (in the case of MBD2) for recruiting the deacetylase core, is unknown (Desai et al., 2015). The localization of full-length MBD3 in NuDe/NuRD is also not known.

Here, we investigated the molecular architecture of the MHR, MHM, and NuDe sub-complexes, using Bayesian integrative modeling with the Integrative Modeling Platform (IMP) (Alber et al., 2007; Rout and Sali, 2019; Russel et al., 2012). This approach allowed us to combine data from multiple sources, including SEC-MALLS, DIA-MS, XLMS, negative-stain EM, X-ray crystallography, NMR spectroscopy experiments, secondary structure and homology predictions, and stereochemistry considerations (Alqarni et al., 2014; Connelly et al., 2006; Cramer et al., 2014; Gnanapragasam et al., 2011; Low et al., 2020; Millard et al., 2016, 2013) to obtain integrative structures with precisions of 27 Å (MHR), 28 Å (MHM), and 25 Å (NuDe). These structures were corroborated by independent crosslinks, cryo-EM maps, biochemical assays, known cancer-associated mutations, and structure predictions from AlphaFold (Desai et al., 2015; Forbes et al., 2006; Millard et al., 2020; Pflum et al., 2001; Zhang et al., 1999). The integrative approach facilitated the modeling of proteins with significant regions of unknown structure, such as full-length MBD3. Our models indicate that MBD3 can potentially localize to two different sites in NuRD, suggesting a mechanism by which MBD3, in the presence of GATAD2A, asymmetrically bridges the deacetylase and chromatin-remodeling modules. Finally, our models enable us to compare the structure of the MHR complex in the presence and absence of MBD3 and GATAD2A. We show that, while the MHR complex alone is relatively dynamic, the presence of MBD3 and GATAD2A makes it less dynamic.

## Results

### Integrative modeling workflow

The modeled NuRD subunits, their domains, their representations, and the number of copies in the modeled complexes are shown in Fig. 1. The representative paralogues used for all calculations were MTA1, HDAC1, RBBP4, MBD3 and GATAD2A (Fig. 1A). The stoichiometry of the modeled proteins was informed by previous DIA-MS and SEC-MALLS experiments (Fig.1B) (Low et al., 2020). The integrative modeling of the MHR, MHM, and NuDe complexes proceeded in four stages (Fig. 2, Material and Methods) (Alber et al., 2007; Rout and Sali, 2019; Russel et al., 2012).

**Fig. 1.**
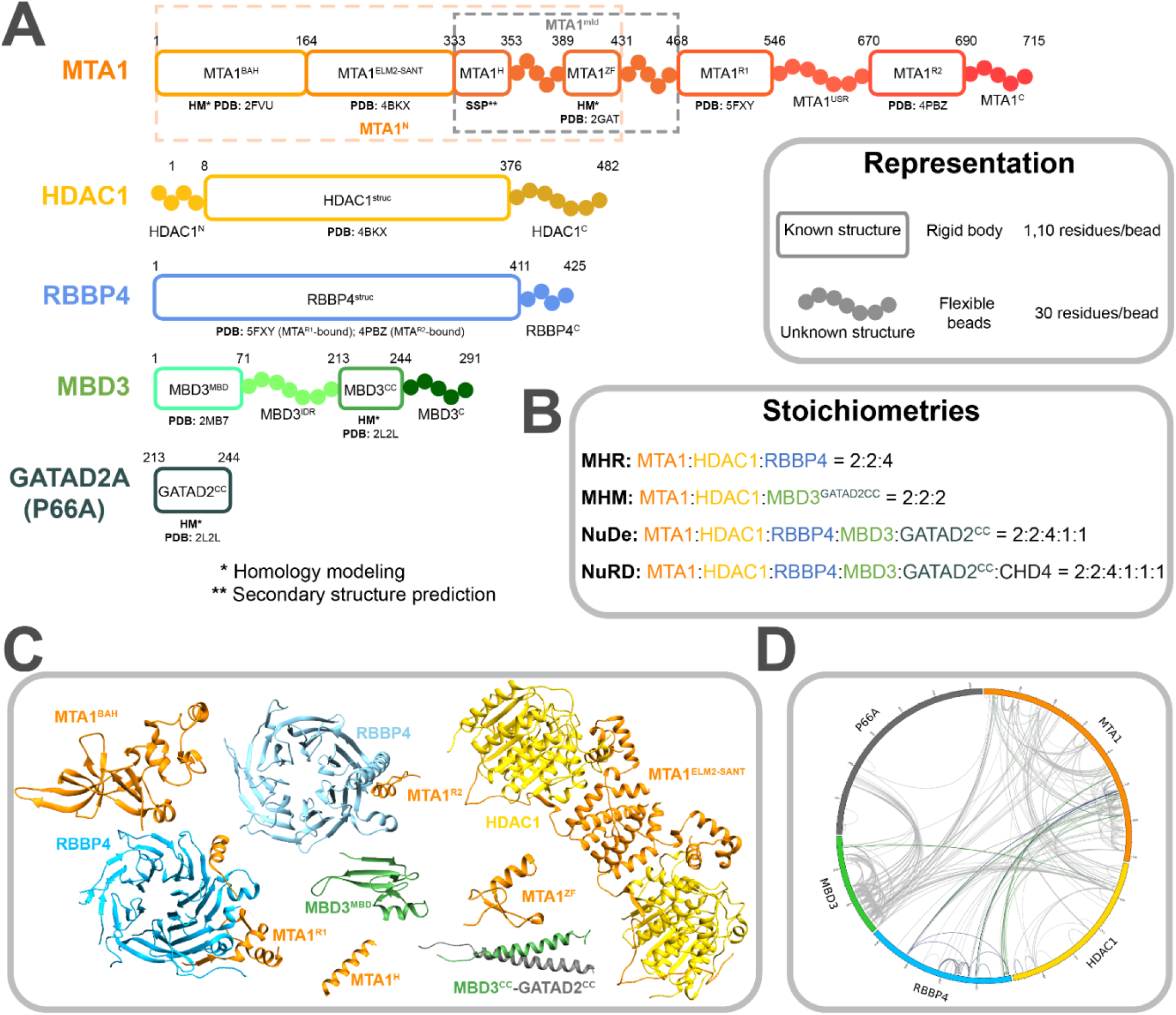
Subunits in NuRD sub-complexes. A. Sequences and isoforms of NuRD subunits modelled in this study are shown and domains are labelled. Only one paralogue of each subunit is shown. Domains are shown in progressively dark shades along the sequence for MTA1, HDAC1, and MBD3. Regions for which atomic structures exist, or can be predicted, are represented by rectangles whereas regions without known structure are represented by beads. PDB IDs are shown for existing subunit structures and templates of homology models. Orange and grey dashed rectangles represent MTA1^N^ and MTA1^mid^ respectively. Numbering is for the human proteins. B. Stoichiometries of modeled sub-complexes and the endogenous NuRD complex. C. Previously published atomic structures that were used for modeling. PDB codes are given in A. D. Crosslinks used in this study, represented as a CIRCOS (http://cx-circos.net/) plot. The grey, blue, and green lines represent all the BS3/DSS, ADH and DMTMM crosslinks respectively.

**Fig. 2.**
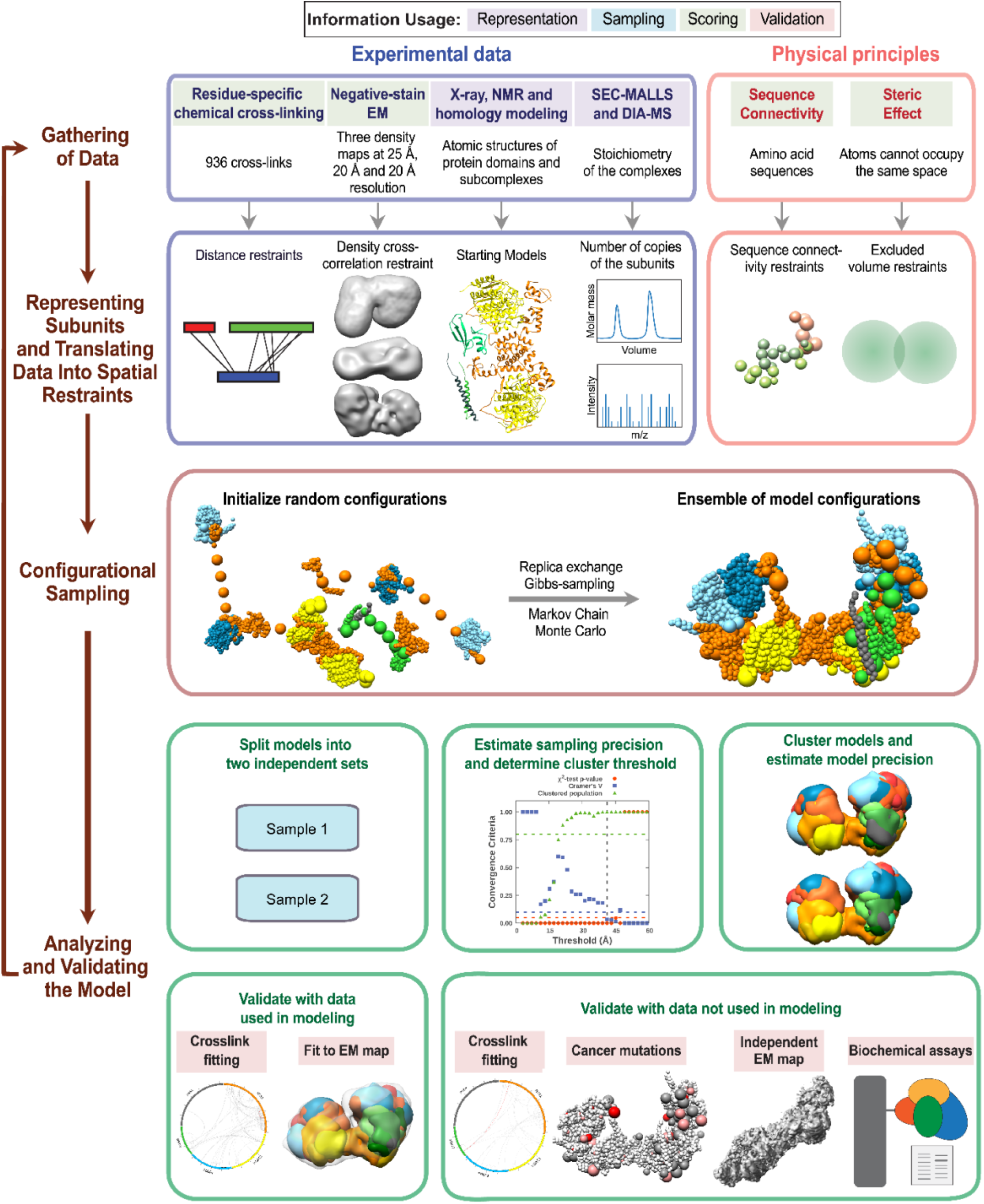
Integrative structure determination of NuRD sub-complexes. Schematic describing the workflow for integrative structure determination of NuRD sub-complexes. The first row describes the input information. The second row details how data are used to encode spatial restraints. The third row describes the sampling method, and the last two rows illustrate the analysis and validation protocol. The background colors of the input information show the stage of modeling in which the information is used, as shown in the legend at the top.

We first represented each protein as a series of beads of sizes that depend on the degree of knowledge of the structure (which can vary throughout the sequence). Protein domains with known atomic structures (such as the MTA1-HDAC1 dimer) were represented at 1 and 10 residues per bead and modeled as rigid bodies, whereas domains without known structure (such as the MBD3^IDR^) were coarse-grained at 30 residues per bead and modeled as flexible strings of beads (Fig.1A). Data from chemical crosslinking combined with mass spectrometry (XL-MS) were used to restrain the distance between crosslinked residues (Fig. 1D). Negative-stain EM maps were used to restrain the shape of the complexes (Fig. S1) (Low et al., 2020).

The simulations started with randomized configurations for the rigid bodies and flexible beads. Over 40 million models per complex were sampled using a Monte Carlo approach (Replica Exchange Gibbs Sampling MCMC; described in Materials and Methods). The models were scored based on agreement with XL-MS and EM data, together with additional stereochemical restraints such as connectivity and excluded volume. For each complex, about 20,000 models that sufficiently satisfied the input information were selected for further analysis (Saltzberg et al., 2021; Viswanath et al., 2017b).

These models were clustered based on structural similarity and the precision of the clusters was estimated (Fig. S2-S4) (Saltzberg et al., 2019, 2021; Viswanath et al., 2017b). The quality of the models was assessed by the fit to input data (Fig. S5-S7), as well as data not used in modeling, such as an independent, published crosslinking dataset (Spruijt et al., 2021), cryo-EM maps (Millard et al., 2020), published biochemical data (Desai et al., 2015; Millard et al., 2020; Pflum et al., 2001; Zhang et al., 1999), human cancer-associated mutations (COSMIC) (Table S1) (Forbes et al., 2006), and predictions from AlphaFold (Jumper et al., 2021). The robustness of the models was also assessed by jack-knifing. 1.3% (MHR), 0% (MHM), and 2% (NuDe) crosslinks in the validation set were violated by these models; they are largely similar to the corresponding models computed with the entire dataset. (Fig. S8, Material and Methods). The resulting integrative models were visualized in two ways – a representative bead model and a localization probability density map – and represented in UCSF Chimera and ChimeraX (Pettersen et al., 2021, 2004). The bead model represents the centroid of the major cluster, whereas the localization probability density map represents all models in the major cluster, by specifying the probability of a voxel (3D volume unit) being occupied by a bead in the set of superposed cluster models.

### MHR

First, to support the integrative modeling of the MHR complex, an *ab initio* 3D EM map for the MHR complex was produced by further analysis of the MHR 2D class averages reported in a previous study (Fig. S1) (Low et al., 2020). Integrative modeling of the 2:2:4 MHR complex produced effectively a single cluster of models (85% of a total of 15200 models) with a model precision of 27 Å; model precision is the average RMSD between the cluster centroid and models in the cluster (Fig. S2). The models fit very well to the input data as measured by the EM and crosslink scores. 98% of the input crosslinks were satisfied within their uncertainty (Fig. S5). An adipic acid dihydrazide (ADH) / bis(sulfosuccinimidyl) suberate or disuccinimidyl suberate (BS3/DSS) / dimethoxy triazinyl methyl-morpholinium chloride (DMTMM) crosslink is violated if the corresponding crosslinked beads are greater than 35 / 35 / 25 Å apart in all models in the cluster. The cross-correlation between the localization probability density map for the models in the major cluster and the input EM map was 0.74, indicating the fit to EM is reasonable but not too high. This could partly be due to unoccupied density in the lobes of the experimental EM map (Fig. 3A).

**Fig. 3.**
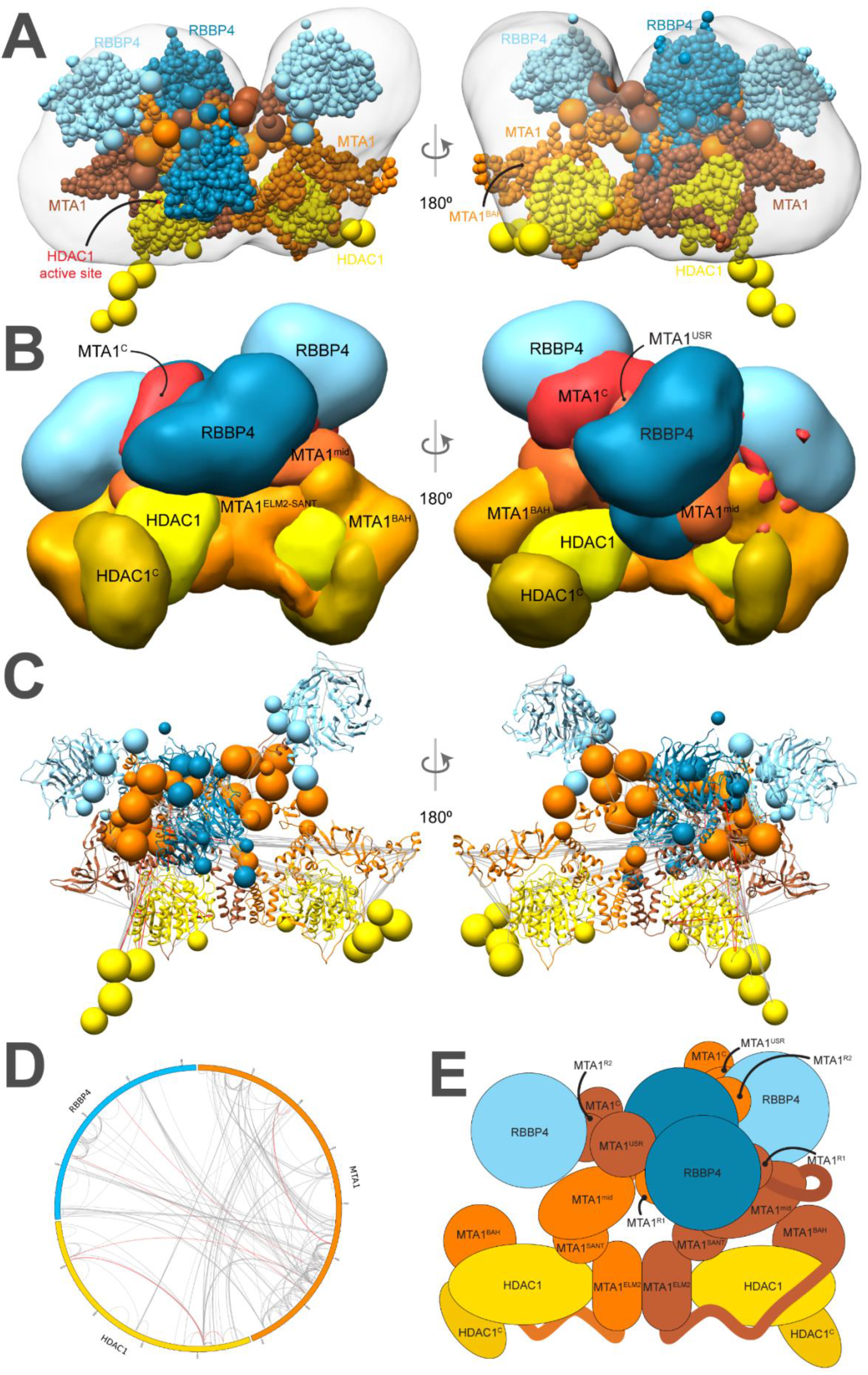
Integrative model of the MTA1-HDAC1-RBBP4 (MHR) complex. A. Representative bead model from the most populated cluster of integrative models for the MHR complex, shown with the MHR EM map. The model is colored by subunit. For MTA1, the two copies are shown in different colors (brown and orange) in panels A and C, to illustrate the crossover. The HDAC1 active site is shown in red. B. Localization probability density maps showing the position of different domains/subunits in the cluster. The map specifies the probability of any volume element being occupied by a domain in the ensemble of superposed models from the cluster. The domain densities are colored according to Fig. 1. These maps are contoured at ∼10% of their respective maximum voxel values. C. Representative bead model from panel A with regions of known structure shown in ribbon representation. D. CX-CIRCOS (http://cx-circos.net/) plot for crosslink satisfaction on the ensemble of MHR models from the major cluster. Grey (red) lines indicate satisfied (violated) crosslinks in panels C and D. E. Schematic representation of the integrative model of the MHR complex. See also Figs. 1, S2, S5.

Surprisingly, the representative bead model from the dominant cluster (cluster centroid model) shows the C-terminal half of the two MTA subunits (MTA1^432-715^) crossing over (brown and orange MTAs, Fig. 3A). Integrative models of the MHR complex created in the absence of the EM map also showed the MTAs crossing over (Fig. S9).

The MTA1^BAH^ domain (MTA1^1-164^) is positioned distal to the MTA1 dimerization interface (MTA1^200-290^, MTA1^dimer^), consistent with its position in an independent EM map (Fig. 3B-3E) (Millard et al., 2020). It is proximal to the HDAC1 active site and therefore might regulate HDAC1 activity (Fig. 3A). This conclusion is consistent with histone deacetylation assays in which MTA1 was shown to modulate HDAC1 deacetylase activity in NuRD (Zhang et al., 1999). Further, for one of the MTAs, the MTA1^BAH^ is located near an RBBP4 (Fig. 3A, Fig. 3B); MTA1^BAH^ proximity to RBBP4 was also indicated in an independent cryo-EM map (Millard et al., 2020). Finally, MTA1^BAH^ is also proximal to the MTA1^mid^ region (MTA1^334-431^) containing the predicted helix (H) and zinc finger regions (ZF) (Fig. 3B, Fig. 3C).

The MTA1^mid^ region is juxtaposed between MTA1^dimer^ and the MTA1^BAH^ domain (Fig. 3B). In contrast, in a previous crosslink-based MHR model (Low et al., 2020), MTA1^mid^ was proximal to the MTA1^BAH^ domain and distal from the MTA1^dimer^. The MTA1 C-terminus (MTA1^C^; *i*.*e*., MTA1^692-715^) shows considerable conformational heterogeneity and is co-located with MTA1^USR^ (MTA1^547-669^), the MTA1 disordered region between the R1 and R2 RBBP4 binding regions (Fig. 3B). Overall, many MTA1 domains in the MHR model, such as MTA1^BAH^ domain, MTA1^mid^, and MTA1^C^, are exposed and could interact with nucleosomal DNA and/or other proteins.

The HDAC1 C-terminus (HDAC1^C^; *i*.*e*., HDAC1^377-482^) interacts with the MTA1^BAH^ domain (Fig. 3B, Fig. 3E). Although it has been shown that the MTA1-HDAC1 dimer can form in the absence of MTA1^BAH^ (Millard et al., 2013), this additional interaction between MTA1 and HDAC1 could be functionally important. Consistent with this possibility, mutations in HDAC1^C^ (Δ391-482, S421A, S423A, E426A) have been shown to disrupt binding to NuRD subunits (Pflum et al., 2001). There are also post-translational modifications in the HDAC1 tail that might modulate its interaction with MTA1 (Pflum et al., 2001; Rathert et al., 2008).

Both the MTA1^R1^-RBBP4 units are located between the two lobes in the EM map, with one complex in the front and the other at the back (dark blue beads and densities, Fig. 3A-3C, Fig. 3E). On the other hand, the MTA1^R2^-RBBP4 complexes are located in separate lobes (light blue beads and densities, Fig. 3A-3C, Fig. 3E). The densities of RBBP4 are spread out, indicating that the localization of these subunits in MHR is imprecise (Fig. 3B). This is consistent with the structural heterogeneity observed in 2D class averages of the MHR EM data (Low et al., 2020). This flexibility is likely necessary to facilitate RBBP4 interactions with transcription factors and histones.

### MHM

Integrative modeling of the 2:2:2 MHM complex resulted in a major cluster containing 99% of the final 28836 models. The model precision was 28 Å and 99% of the input crosslinks were satisfied (Fig. S3, Fig. S6). The cross-correlation between the localization probability density map for the models in the major cluster and the input EM map was 0.9.

Our 2:2:2 MHM model shows two binding sites of MBD3 on the MTA1-HDAC1 dimer (Fig. 4A-4C). Both copies of MBD3^MBD^ localize close to the MTA1^BAH^ domain, which is consistent with the location observed for MBD2^MBD^ in an independent cryo-EM map of a 2:2:1 MTA1:HDAC1:MBD2 complex (Millard et al., 2020) and an independent set of crosslinks (Spruijt et al., 2021) (Fig. 4A-4E). Although there are two MBD3s in our models, a single MBD3^IDR^ localizes to the MTA1 dimerization interface, MTA1^dimer^ (green MBD3, Fig. 4B, Fig. 4E). This localization of MBD3^IDR^ is consistent with its previously predicted localization from a crosslink-based model (Low et al., 2020) and the putative localization of MBD2^IDR^ based on cryo-electron microscopy (Millard et al., 2020). It is also supported by two separate mutagenesis and co-immunoprecipitation studies, one of which showed that MBD2^IDR^ was essential for binding to the MTA1-HDAC1 dimer (Desai et al., 2015), whereas the other showed that MTA1^dimer^ was essential for the interaction with MBD2 (Millard et al., 2020). Finally, the MBD3^CC^-GATAD2^CC^ coiled-coil domain is exposed.

**Fig. 4.**
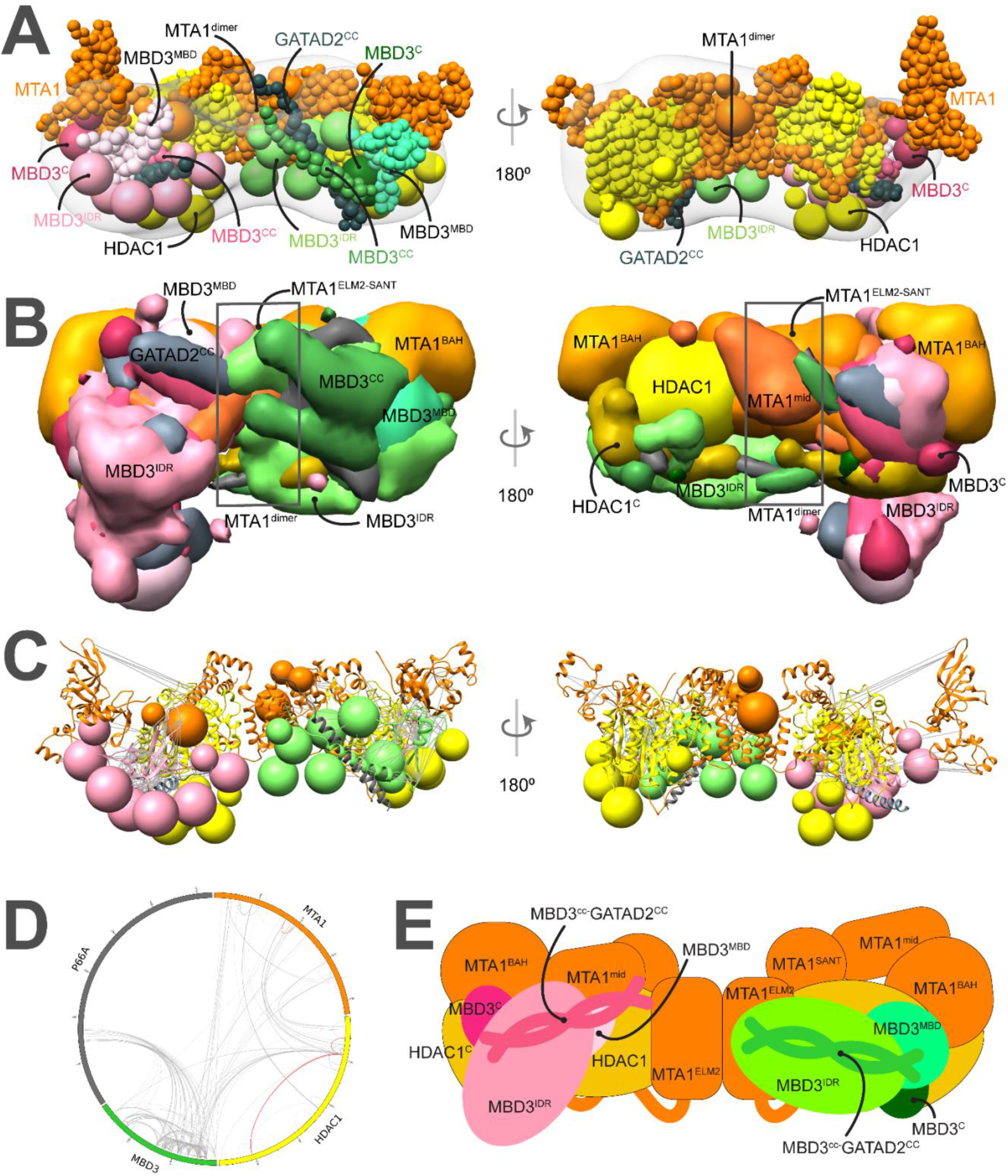
Integrative model of the MTA1^N^-HDAC1-MBD3^GATAD2CC^ (MHM) complex. A. Representative bead model from the major cluster of analyzed integrative models for the MHM complex, with the corresponding EM map (EMD-21382) (Low et al., 2020), colored by subunit. The domains of the two MBD3s are shown in shades of pink and green, respectively. B. Localization probability density maps showing the position of different domains in the ensemble of models from the cluster. The domain densities are colored according to Fig 1. C. The same density maps as B (front view), showing the two MBDs in pink and green, respectively, and illustrating that they localize differently on the MTA1-HDAC1 dimer. The density maps of MTA1^mid^ and GATAD2^cc^ were omitted for clarity. D. The density maps of the two MBD3^IDR^ domains on the MTA1-HDAC1 dimer. Most of the maps are contoured at around 20% of their respective maximum voxel values (except MTA1^165-333^ at 10% and GATAD2^cc^ at 27%). C. Representative bead model from panel A with regions of known structure shown in ribbon representation. D. CX-CIRCOS (http://cx-circos.net/) plot for crosslinks satisfaction on the ensemble of MHM models from the major cluster. Grey (red) lines indicate satisfied (violated) crosslinks in panels C and D. E. Schematic representation of the integrative model of the MHM complex. E. Schematic representation of the integrative model of the MHM complex. Note that MTA1^mid^ in this model corresponds to MTA1^334-431^. See also Figs. 1, S3, S6.

### NuDe

In modeling the NuDe complex, we incorporated only the region of GATAD2A that forms a coiled-coil with MBD3, because of the lack of structural information on GATAD2A and the very small number of XLs involving this subunit. Integrative modeling of the NuDe complex resulted in effectively a single cluster (92% of 19754 models). The model precision was 34 Å and 99% of the input crosslinks were satisfied (Fig. S4, Fig. S7). The cross-correlation between the localization probability density map for the models in the major cluster and the input EM map was 0.9. Further, we validated these models by an independent set of crosslinks on the endogenous NuRD complex and interacting proteins (Spruijt et al., 2021); all 89 crosslinks used in this analysis were satisfied.

In contrast to the MHM model, MBD3 is localized more precisely in NuDe. It is juxtaposed next to the MTA1^BAH^ and MTA1^mid^ domains (Fig. 5A-5E). An independent cryo-EM map of MTA1^1-546^-HDAC1-MBD2-RBBP4 (Millard et al., 2020) and an independent set of crosslinks (Spruijt et al., 2021) also showed that MBD3 was proximal to MTA1^BAH^ and MTA1^dimer^. Similar to the MHM model, MBD3^IDR^ extends towards MTA1^dimer^.

**Fig. 5.**
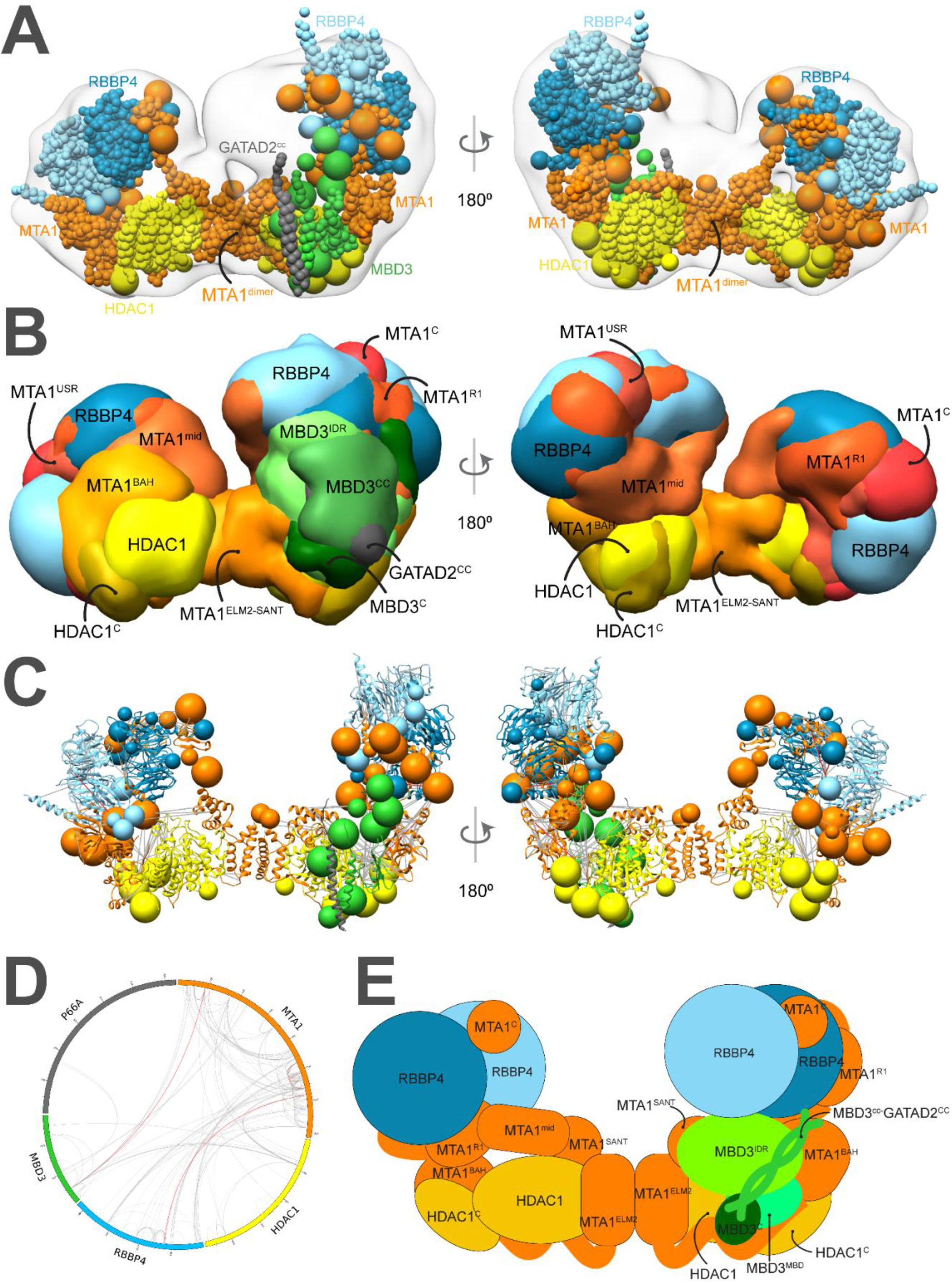
Integrative model of the nucleosome deacetylase (NuDe) complex. A. Representative bead model from the dominant cluster of integrative models for the NuDe complex, with the corresponding EM map (EMD-22904) (Low et al., 2020), colored by subunit. B. Localization probability density maps showing the position of different domains in the ensemble of models from the cluster. The domain densities are colored according to Fig 1. Maps are contoured at ∼10% of their respective maximum voxel values (except GATAD2^CC^ at 20%). C. Representative bead model from panel A with regions of known structure shown in ribbon representation. D. CX-CIRCOS (http://cx-circos.net/) plot for crosslinks satisfaction on the ensemble of NuDe models from the major cluster. Grey (red) lines indicate satisfied (violated) crosslinks in panels C and D. E. Schematic representation of the integrative model of the NuDe complex. See also Figs. 1, S4, S7, S10.

From protein-protein distance maps of the cluster, HDAC1^60-100^ and MTA1^BAH^ are most proximal to MBD3 (Fig. S10A, S10B). The MBD3^CC^-GATAD2^CC^ coiled-coil is exposed. The MBD3^MBD^ domain is buried, consistent with the failure of MBD3 to bind DNA in NuRD noted in immunoprecipitation experiments (Fig. 5A-5E) (Zhang et al., 1999). Interestingly, several nucleosome-interacting domains such as MTA1^BAH^ and MTA1^ZF^ are co-localized in the NuDe model (Fig. 5A-5E).

Similar to the MHR models, the HDAC1^C^ domain is proximal to MTA1^BAH^ (Fig. S10C). In contrast to the MHR models, which showed crossover of MTAs, the two MTAs are well-separated in NuDe (Fig. 5A-5E). The localization of RBBPs is also more precise in NuDe than in MHR (Fig. 3B, Fig. 5B).

### Mapping COSMIC mutations

We next consulted the COSMIC (Catalogue of Somatic Mutations in Cancer) database for somatic, confirmed pathogenic, point mutations of the NuRD subunits, MTA1, HDAC1, RBBP4, and MBD3 (Forbes et al., 2006). In total, 356 point mutations were identified and mapped onto the cluster of NuDe integrative models (Methods, 5.1 COSMIC data analysis). Analysis of these mutations revealed that 24% of residues with three or more reported COSMIC mutations mapped to exposed regions that are known to bind to nucleosomes and transcription factors, such as the HDAC1 active site and RBBP4 H3 interaction site (Fig. 6, Fig. S11, Table S1). This number is significantly higher than the number expected based on random chance (9%) (Materials and Methods). However, 44% of the residues with three or more reported COSMIC mutations, in comparison to 41% in the random set, mapped to subunit interfaces in NuDe. This suggests that protein-protein interfaces within NuRD are not over-represented in the COSMIC database, whereas mutations at exposed surfaces are. This perhaps indicates that interactions of NuRD subunits with other macromolecules, such as nucleosomes and/or transcription factors, are crucial for the function of the complex. Therefore, mutations that impair binding of NuRD to its binding partners could contribute to the pathogenesis of the disease.

**Fig. 6.**
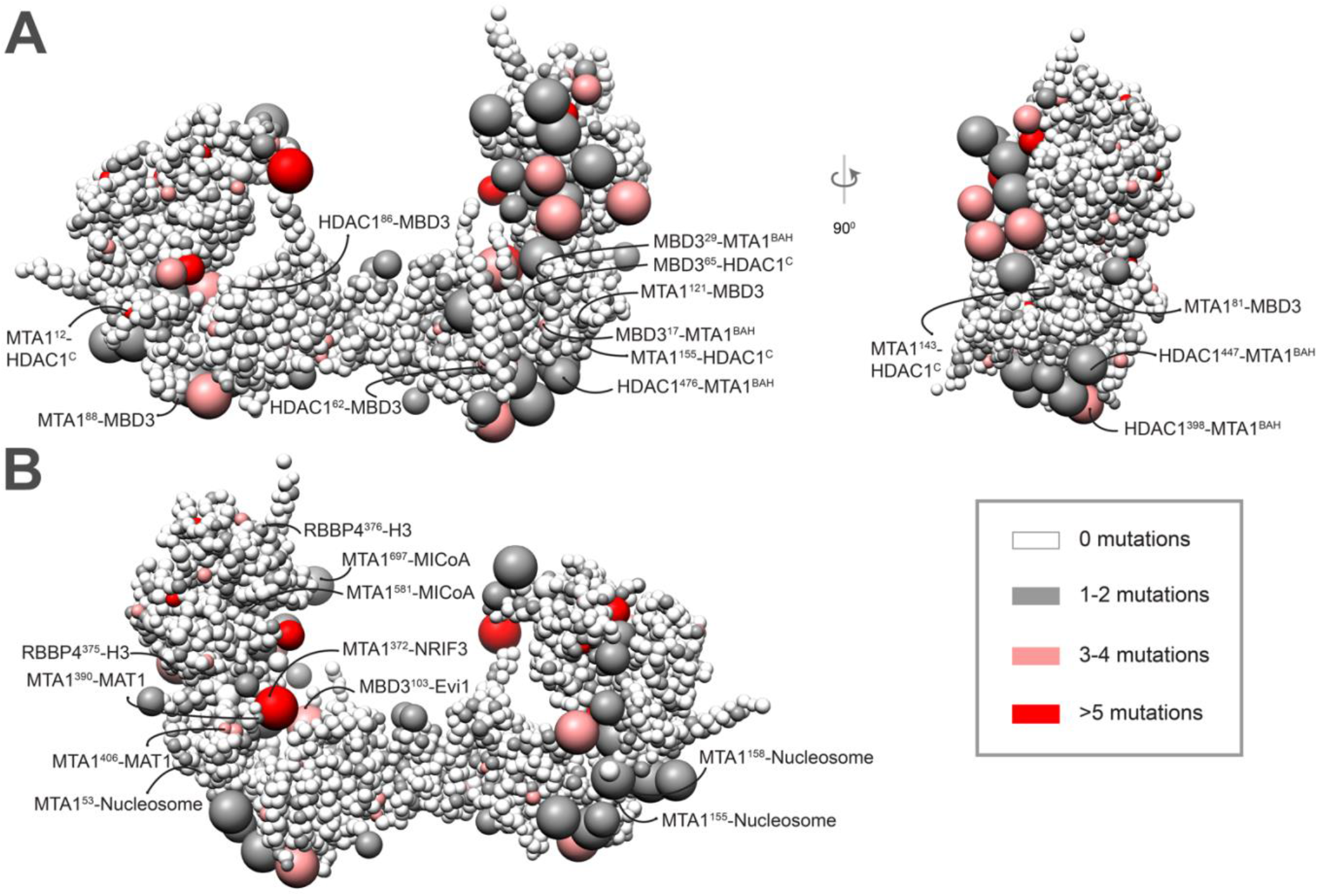
COSMIC mutations mapped onto the NuDe integrative model. Somatic pathogenic point mutations from the COSMIC database (Forbes et al., 2006) mapped onto the representative bead model of the NuDe complex (Fig. 5A). A. Mutations of residues that map to previously undescribed protein-protein interfaces within our model. B. Mutations on residues that map to exposed binding sites between modeled proteins and known binding partners. A bead is colored according to the maximum number of mutations on any residue in the bead, according to the legend. Representative mutations are labeled in both A. and B. See also Table S1 and Fig. S12.

Moreover, of the 24% of mutations that map to exposed regions, half map to regions of unknown structure (regions for which no experimental structure or reliable model is available), such as MTA1^USR^ and MBD3^IDR^ (Fig. 6, Table S1). The functional significance of these mutations is therefore difficult to predict but could indicate that these regions of unknown structure also have important roles in protein stability, regulating interactions with binding partners of NuRD, or interactions between NuRD subunits. An important additional consideration for all these disease-causing mutations is that many of the NuRD subunits function in cellular contexts independent of other NuRD subunits, and so in some cases, these mutations may be rationalized in the context of other functional roles.

### Docking the nucleosome

We next attempted to dock the CHD4-nucleosome structure (Farnung et al., 2020) into the cleft in the NuDe structure between the MTA1 C-terminal arms (Fig. 7). Although there are limitations to this docking, this positioning of the nucleosome indicates its size complementarity to the integrative model, further corroborating the latter. This placement allows for the histone tails to be located towards the HDAC1 active site. It also accommodates the known interactions between the RBBPs and the histone H3 (Fig. 7). The partial CHD4 structure is exposed. MTA1^mid^, which contains the zinc finger, can also potentially interact with the nucleosome in this position. Finally, MBD3 does not interact with the nucleosome, since MBD3^MBD^ is buried in our model of NuDe (Fig. 5B), consistent with MBD3 in NuRD failing to bind DNA in immunoprecipitation experiments (Zhang et al., 1999).

**Fig. 7.**
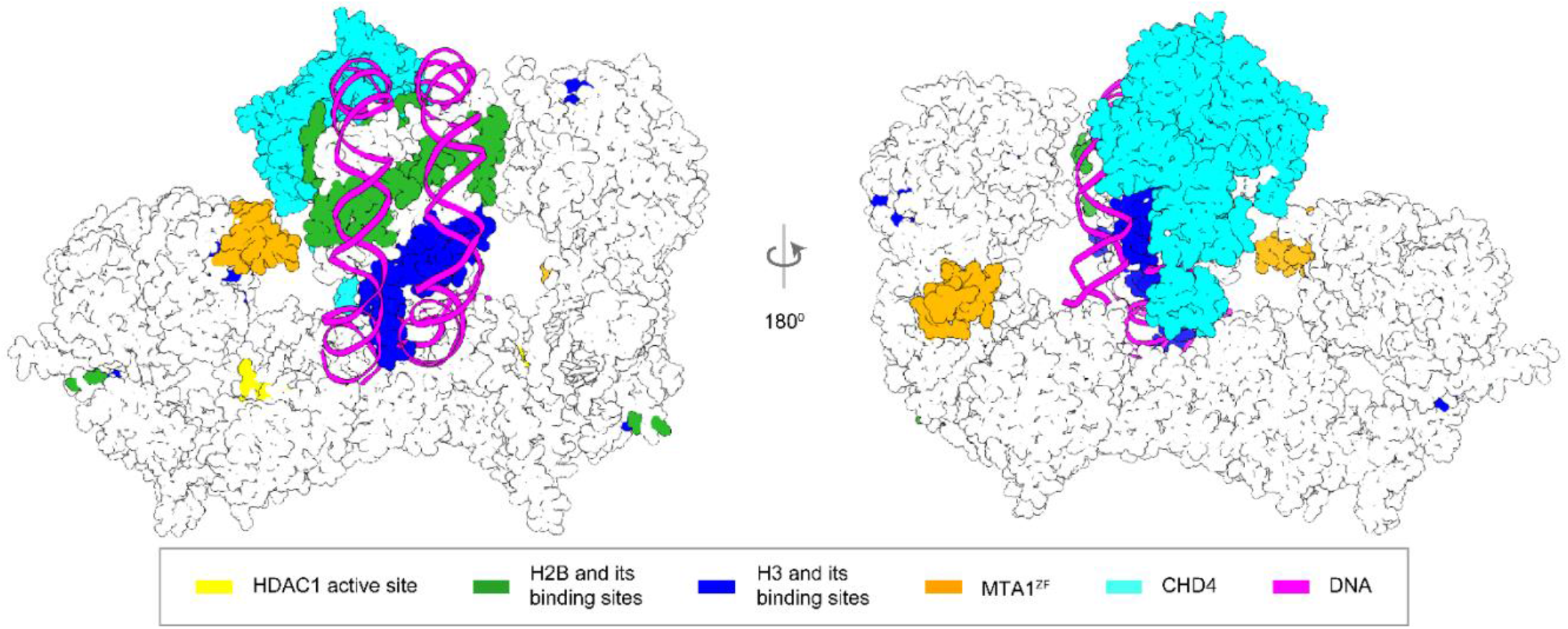
Integrative model of NuDe complex with the nucleosome. The CHD4-nucleosome structure (Farnung et al., 2020) is placed in the cleft of the NuDe integrative model. The regions with known atomic structure are shown in the NuDe integrative model from Fig. 5A. CHD4, histones, DNA and the corresponding NuDe subunit residues they are proposed to bind to, are depicted in the same color, as given by the legend.

### Comparison to structure predictions by AlphaFold-Multimer

We compared our models to models predicted by AlphaFold-Multimer for a 2:2:1 MTA1^1-350^-HDAC1^1-376^-MBD3^1-291^-GATAD2A^213-244^ complex (Fig. S12) (Evans et al., 2022). In the top AlphaFold prediction, the MTA1-HDAC1 dimer closely resembles the corresponding crystal structure (Millard et al., 2013), as expected, given that AlphaFold is trained on the PDB. The localization of MTA1^BAH^, MTA1^H^, and MBD3 are broadly consistent with our integrative model and the cryo-EM map from an independent study (Millard 2021). MBD3^MBD^ is proximal to MTA1^BAH^, as predicted in our model. MBD3^IDR^ is near the MTA1 dimerization interface. Specifically, the N-terminal part of the MBD3^IDR^ (MBD3^125-175^) winds an irregular path towards the MTA1^dimer^, whereas the C-terminal part of the MBD3^IDR^ (MBD3^125-175^), which is predicted to be ordered, forms a compact structure at the MTA1^dimer^ (Fig. S12). The MBD3^CC^-GATAD2A coiled-coil sits diagonally across the MTA1 dimerization interface. This prediction, however, has several clashes (a phenomenon that we and others have observed in other predictions made using AlphaFold multimer, unpublished) and violates some of the input crosslinks. Thus, 94%, 87%, 94%, 87%, 61%, and 61% of the HDAC1-HDAC1, MTA1-MTA1, HDAC1-MTA1, MBD3-MTA1, MBD3-MBD3, and HDAC1-MBD3 crosslinks, respectively, were satisfied by the top AlphaFold model.

## Discussion

Here, we obtained structural models of the MTA1-HDAC1-RBBP4 (MHR), MTA1^N^-HDAC1-MBD3 (MHM), and MTA1-HDAC1-RBBP4-MBD3-GATAD2A (NuDe) complexes using Bayesian integrative modeling with IMP. Our approach has the several advantages over other modeling strategies. First, models generated by IMP can incorporate full-length protein sequences, including regions that are predicted to be intrinsically disordered and/or might only form structure once assembled in a complex. IMP allows for a multi-scale representation, with regions of known atomic structure and unknown structure represented at higher and lower resolutions, respectively. This feature facilitated the modeling of NuRD proteins with significant regions of unknown structure, such as full-length MBD3, for the first time. Second, the Bayesian inference framework in IMP allows us to combine data from several experiments at multiple resolutions, *e*.*g*., negative-stain EM and XLMS, by considering the data uncertainty, and without using arbitrary weights, making the sampling more objective and accurate (Alber et al., 2007; Rieping et al., 2005; Rout and Sali, 2019; Schneidman-Duhovny et al., 2014). Finally, the output of IMP is an ensemble of models consistent with the data, instead of a single model. This allows us to obtain precise uncertainty bounds on the structure as a measure of model precision (Saltzberg et al., 2019, 2021; Viswanath et al., 2017b; Webb et al., 2018).

### Comparison between IMP and AlphaFold

Integrative models are computed by satisfying spatial restraints based on information specific to a given system, whereas AI-based methods such as AlphaFold predict structures relying heavily on features learnt from general databases of protein structures. The advantages of the latter methods are that they are fast, easy to use, and generate structures at atomic resolution. However, the predicted structures may not fit experimental data for a given system. For example, for our complex, the top prediction from AlphaFold violated a significant number of crosslinks. In contrast, integrative structure determination by IMP produces an ensemble of models consistent with the input information, instead of a single structure. In addition, rigorous validation of the integrative models based on fit to input information, jack-knifing, and consistency with information not used in modeling, is an essential part of the integrative approach.

### NuDe complex is more ordered than MHR

A comparison of MTA1 and RBBP4 in the MHR and NuDe models suggests that these subunits are more conformationally heterogenous in MHR, as shown by the broader localization probability densities for the C-terminal half of MTA1 and RBBPs in MHR (volume enclosed by the corresponding maps = 1400 nm^3^) compared to NuDe (volume enclosed = 1300nm^3^). Also, the cross-correlation of the MHR localization probability density to the corresponding EM map is lower than that of NuDe, indicating higher heterogeneity for the former. This result suggests that dynamics in the MHR sub-complex might be damped in the presence of MBD3-GATAD2A.

### MBD3^IDR^ – MTA1^dimer^ interaction

In our MHM models, one MBD3^IDR^ is near the MTA1^dimer^, consistent with the previously predicted localization of MBD3^IDR^ based on chemical crosslinks (Low et al., 2020) and MBD2^IDR^ based on a cryo-electron density map (Fig. 4C-4E) (Millard et al., 2020). Two separate mutagenesis and co-immunoprecipitation studies have shown that the MBD^IDR^ and the MTA1 dimerization interface are each essential for MBD2 interaction with the MTA1-HDAC1 dimer (Desai et al., 2015; Millard et al., 2020). Despite the corresponding region of MBD2 being disordered in solution when the protein is in isolation (Desai et al., 2015), MBD3^125-175^ is predicted to be ordered based on PONDR^®^ analysis (Fig. S13) (http://www.pondr.com) (Romero et al., 2001, 1997). Because this region is well conserved across species (Cramer et al., 2017), it is likely that it becomes ordered upon binding, similar to the region of MTA1 that winds irregularly across the surface of HDAC1 (MTA1^165-226^). Further, the crosslinks between MBD3^IDR^ and MTA1 involve a loop (MTA1^229-236^) of the MTA1^dimer^ that is not visible in the MTA1-HDAC1 crystal structure. It is possible that this region of MTA1 may also become ordered upon binding MBD3. It is also worth noting that, unlike the NuDe and MHR complexes, MHM has not been observed in cells or in material purified from cells. MHM was created as part of a reductionist strategy towards understanding NuRD assembly

### Model of MBD3 binding in NuRD

The stoichiometry of MBD3 in NuRD is intriguing. The MHM complex has two copies of MBD3, though it is likely that this is not a physiologically relevant complex, whereas a single MBD3 is seen in the NuDe and NuRD complexes (Low et al., 2020). Based on our integrative models, we propose a two-state mechanism to explain the asymmetric binding of MBD3 in NuRD (Fig. 8).

**Fig. 8.**
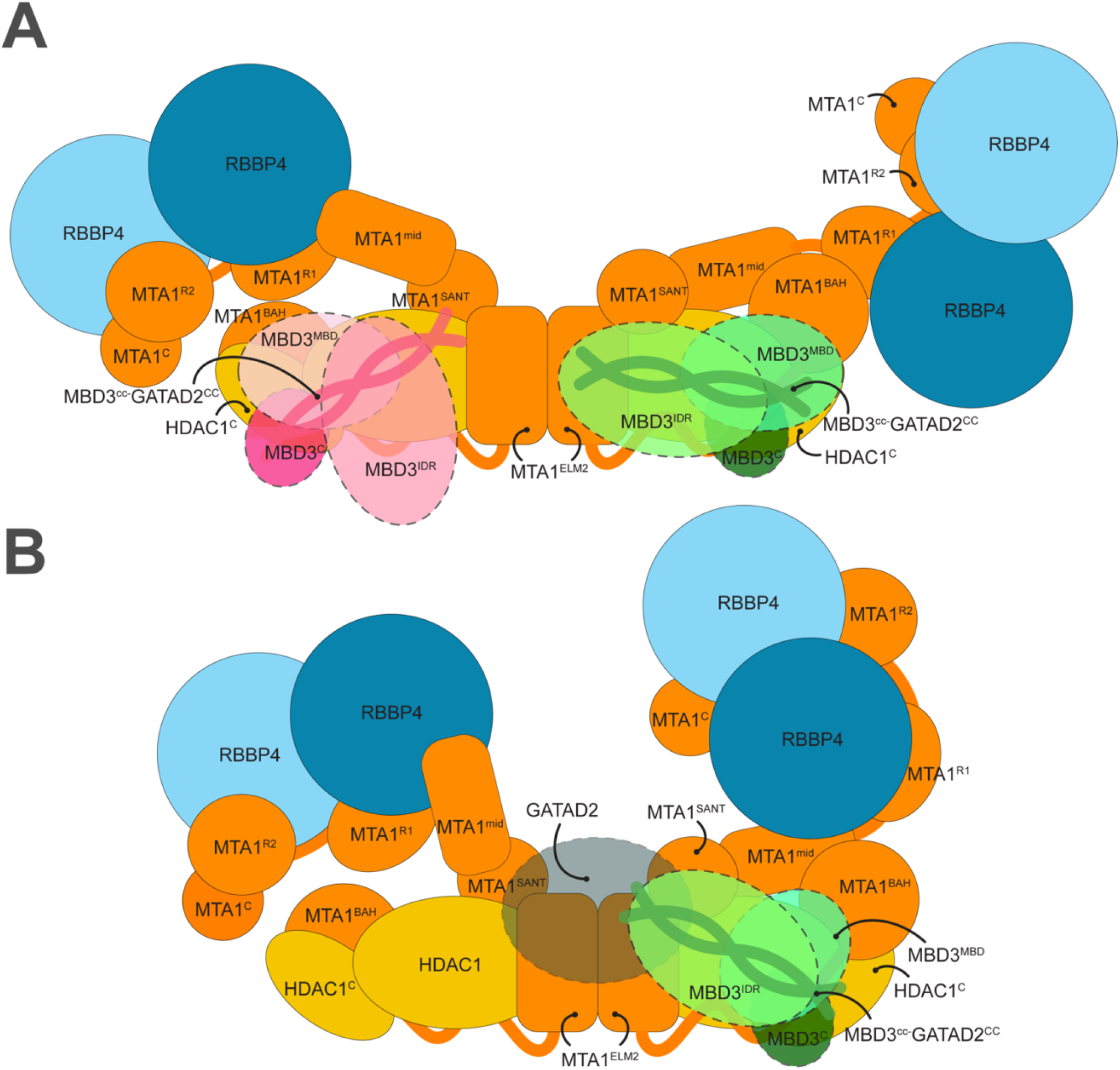
Model of MBD3 binding in NuRD. The figure shows two states of MBD3 in NuRD. A. In the first state, the MTA1 dimerization interface is accessible for MBD3^IDR^ to bind. B. In the second state, upon binding, MBD3 recruits GATAD2A and the chromatin remodeling module and shifts to one end of the MTA1-HDAC1 dimer. GATAD2A localizes near MTA1dimer, precluding a second MBD3 from binding to it.

In the first state (Fig. 8A), the C-terminal arms of MTA1 in MHR are heterogenous and adopt a range of configurations including an extended, open configuration (Millard et al., 2020) and crossed-over configuration (Fig. 3, MHR models). Functionally, this flexibility could play a role in facilitating interactions with histones as well as the transcriptional regulators such as PWWP2A that can recruit MHR to target sites (Link et al., 2018). In the open state, the MTA1 dimerization interface is accessible for MBD3^IDR^ to bind. This MBD3^IDR^-MTA1^dimer^ interaction is critical for MBD recruitment to the deacetylase module (Fig. 4, MHM models) (Desai et al., 2015; Millard et al., 2020). Although there are two MBD3 binding sites on the MTA1-HDAC1 dimer, only one interacts with the MTA1 dimer. This is probably the MBD3 that is present in the physiological NuDe/NuRD complex.

In the second state (Fig. 8B), upon binding to MTA1^dimer^, MBD3 recruits GATAD2A and the chromatin remodeling module, and shifts to one end of the MTA1-HDAC1 dimer (Fig. 5, NuDe models). In this state, GATAD2A localizes near MTA1^dimer^, precluding a second MBD3 from binding to it. Although we did not model full GATAD2A in NuDe due to unavailability of structures and crosslinks involving the protein, the proximity of CHD4, and hence GATAD2A, to the MTA1^dimer^ in our coarse nucleosome docking supports this idea (Fig. 7). This possibly explains how GATAD2A introduces asymmetry of MBD3 binding in NuRD. Moreover, upon binding the chromatin remodeling module, the C-terminal arms of MTA1 with the RBBPs are less heterogenous and adopt a closed configuration (Fig. 3B, Fig. 5B). This reduced flexibility of MTA1^C^-RBBP in NuDe may reflect a functional distinction between the MHR and NuDe sub-complexes, the nature of which is not currently clear.

In the second state, the MBD3^MBD^ is buried and fails to bind DNA, also noted in previous experiments (Zhang et al., 1999). MBD3 is less effective at distinguishing methylated and unmethylated DNA compared to other MBD paralogs (Liu et al., 2019). Considering these facts, it is possible that the major role of MBD3 here is to connect its two enzymatic modules rather than to recruit NuRD to DNA, methylated or otherwise. Experiments in mouse ES cells showed that MBD3 deletion resulted in loss of integrity of NuRD in these cells, supporting its role in NuRD assembly (Kaji et al., 2006).

The novel NuRD protein interfaces predicted by our model need to be confirmed by future experiments. High-resolution structures of regions such as MBD3^IDR^ will delineate their roles in NuRD, although they will continue to be a challenge to characterize by empirical methods. Ultimately, a complete atomic characterization of the NuRD complex will aid in understanding NuRD-mediated regulation of gene expression.

## Materials and Methods

### Integrative modeling

The integrative structure determination of the NuRD sub-complexes proceeded through four stages (Fig. 2) (Alber et al., 2007; Rout and Sali, 2019; Russel et al., 2012). The modeling protocol (i.e., stages 2, 3, and 4) was scripted using the Python Modeling Interface (PMI) package, a library for modeling macromolecular complexes based on open-source Integrative Modeling Platform (IMP) package, version 2.13.0 (https://integrativemodeling.org) (Russel et al., 2012). The current procedure is an updated version of previously described protocols (Ganesan et al., 2020; Gutierrez et al., 2020; Kim et al., 2018; Saltzberg et al., 2019, 2021; Viswanath et al., 2017a; Webb et al., 2018). Files containing the input data, scripts, and output results are publicly available at https://github.com/isblab/nurd.

### Stage 1: Gathering data

The stoichiometry and isoforms of subunits were based on DIA-MS and SEC-MALLS experiments (Fig. 1) (Low et al., 2020). Known atomic structures were used for the MTA1-HDAC1 dimer, MTA1^R1^ and MTA1^R2^ domains in complex with RBBP4, and MBD domain of MBD3 (Fig. 1) (Alqarni et al., 2014; Cramer et al., 2014; Millard et al., 2016, 2013). The MTA1^BAH^ domain, MTA1^H^, MTA1^ZF^, and MBD3^CC^-GATAD2A^CC^ structures were homology-modeled based on the structures of related templates (Fig. 1A) (Connelly et al., 2006; Gnanapragasam et al., 2011; Tjandra et al., 1997).

The shapes of the complexes were based on 3D negative-stain EM maps; MHR: to be deposited (24.56 Å), MHM: EMD-21382 (20 Å), and NuDe: EMD-22904 (20 Å) (Low et al., 2020). The negative-stained EM map for the MHR complex was produced by further analysis of data reported in a previous study (Fig. S1) (Low et al., 2020). 25,155 particle images were subjected to multiple rounds of 2D classification in CryoSparc (Punjani et al., 2017), following which an *ab initio* 3D reconstruction was obtained and refined by homogenous 3D refinement. The final map was produced from 13,299 particles and had an estimated resolution of ∼25 Å according to the FSC0.143 criterion.

Chemical crosslinks informed the relative localization of the NuRD subunits. For modeling the MHR complex, 387 BS3/DSS (bis(sulfosuccinimidyl)suberate and disuccinimidyl suberate), 34 DMTMM (dimethoxy triazinyl methyl-morpholinium chloride), and 19 ADH (adipic acid dihydrazide) crosslinks were used (Low et al., 2020). For MHM (NuDe), 539 (312) BS3/DSS, 0 (40) DMTMM and 0 (19) ADH crosslinks were used for modeling.

The models were validated by independent EM maps (Millard et al., 2020), biochemical assays (Desai et al., 2015; Pflum et al., 2001; Zhang et al., 1999), and human cancer-associated mutations on NuRD proteins (Forbes et al., 2006).

### Stage 2: Representing the system and translating data into spatial restraints

The stoichiometry and representation of subunits is shown (Fig. 1). The domains with known atomic structures were represented in a multi-scale manner with 1 and 10 residues per bead to maximize computational efficiency. These domains were modeled as rigid bodies where the relative distances between beads is constrained during sampling. In contrast, domains without known structure were coarse-grained at 30 residues per bead and modeled as flexible strings of beads.

We next encoded the spatial restraints into a scoring function based on the information gathered in Stage 1, as follows:

#### (1) Crosslink restraints

The Bayesian crosslinks restraint (Rieping et al., 2005) was used to restrain the distances spanned by the crosslinked residues (Shi et al., 2014). The restraint accounts for ambiguity (multiple copies of a subunit) via a compound likelihood term that considers multiple residue pairs assigned to an individual crosslink (Shi et al., 2014). Intra-subunit crosslinks are also considered ambiguous when there are multiple copies of a subunit.

#### (2) EM restraints

The Bayesian EM density restraint was used to restrain the shape of the modeled complexes and was based on the cross-correlation between the Gaussian Mixture Model (GMM) representations of the NuRD subunits and the GMM representation of the corresponding negative-stain EM density maps (Bonomi et al., 2019).

#### (3) Excluded volume restraints

The excluded volume restraints were applied to each bead, using the statistical relationship between the volume and the number of residues that it covered (Alber et al., 2007).

#### (4) Sequence connectivity restraints

We applied the sequence connectivity restraints, using a harmonic upper distance bound on the distance between consecutive beads in a subunit, with a threshold distance equal to twice the sum of the radii of the two connected beads. The bead radius was calculated from the excluded volume of the corresponding bead, assuming standard protein density (Shi et al., 2014).

### Stage 3: Structural sampling to produce an ensemble of structures that satisfies the restraints

We aimed to maximize the precision at which the sampling of good-scoring solutions was exhaustive (Stage 4). The sampling runs relied on Gibbs sampling, based on the Replica Exchange Monte Carlo algorithm (Saltzberg et al., 2019, 2021). The positions of the rigid bodies (domains with known structure) and flexible beads (domains with unknown structure) were sampled.

The initial positions of the flexible beads and rigid bodies in all complexes were randomized, with one exception. For MHR, we were able to unambiguously dock the structure of the MTA1-HDAC1 core in the EM map, with the help of the previous EM map (EMD-3399) (Millard et al., 2016). Hence, the position of the corresponding rigid body was fixed throughout.

The Monte Carlo moves included random translations of individual beads in the flexible segments and rigid bodies (around 3.7 Å and 1.3 Å respectively). A model was saved every 10 Gibbs sampling steps, each consisting of a cycle of Monte Carlo steps that moved every bead and rigid body once.

The sampling produced a total of 40 million MHR, 48 million MHM, and 80 million NuDe integrative models.

### Stage 4: Analysing and validating the ensemble of structures and data

The sampled models were analysed to assess sampling exhaustiveness and estimate the precision of the structure, its consistency with input data and consistency with data not used in modeling. The structure was further validated by experiments based on the predictions from the models. We used the analysis and validation protocol published earlier (Rout and Sali, 2019; Saltzberg et al., 2019, 2021; Viswanath et al., 2017b). Assessment began with a test of the thoroughness of structural sampling, including structural clustering of the models, estimating model precision, and visualizing the variability in the ensemble of structures using localization probability density maps (Viswanath et al., 2017b). The precision of a domain refers to its positional variation in an ensemble of superposed models. It can also be visualized by the localization probability density map for the domain. A localization probability density map specifies the probability of a voxel (3D volume unit) being occupied by a bead in a set of superposed models. The models and densities were visualized with UCSF Chimera and ChimeraX (Pettersen et al., 2021, 2004).

#### (1) Determining good-scoring models

Starting from the millions of sampled models, first, we selected models obtained after score equilibration and clustered them based on the restraint scores (Saltzberg et al., 2021). For further analysis, we considered 15200 MHR, 28836 MHM, and 19754 NuDe good-scoring models that satisfy the data restraints sufficiently well.

#### (2) Clustering and structure precision

We next assessed the sampling exhaustiveness and performed structural clustering (Saltzberg et al., 2019, 2021; Viswanath et al., 2017b). Integrative structure determination resulted in effectively a single cluster for all complexes, at a precision of 27 Å (MHR), 28 Å (MHM), and 34 Å (NuDe). The cluster precision is the bead RMSD from the cluster centroid model averaged over all models in the cluster (Viswanath et al., 2017b).

#### (3) Fit to input information

The dominant clusters from each modeled NuRD sub-complex satisfied over 95% of all the BS3/DSS, ADH, and DMTMM crosslinks used; a crosslink is satisfied by a cluster of models if the corresponding Cα-Cα distance in any model in the cluster is less than 35Å, 35Å, 25Å for BS3/DSS, ADH, and DMTMM crosslinks respectively. The agreement between the models and the corresponding EM maps was computed by calculating the cross-correlation of the combined localization probability densities of all subunits for the major cluster with the experimental EM map using the *fitmap* tool in UCSF Chimera (Fig. 3-5) (Pettersen et al., 2004). The remainder of the restraints are harmonic, with a specified standard deviation. The cluster generally satisfied the excluded volume and sequence connectivity restraints. A restraint is satisfied by a cluster of models if the restrained distance in any model in the cluster (considering restraint ambiguity) is violated by less than 3 standard deviations, specified for the restraint. Most of the violations are small, and can be rationalized by local structural fluctuations, coarse-grained representation of the model, and/or finite structural sampling.

#### (4) Jack-knifing

The robustness of the models was assessed by jack-knifing, *i*.*e*., generating models using a subset of the input crosslinks. For each modeled sub-complex, we generated models based on a randomly selected subset consisting of 80% of the BS3/DSS crosslinks and the corresponding EM maps, and used the remaining 20% of the crosslinks for validation. Our analysis showed that 1/79 (MHR), 0/111 (MHM) and 1/64 (NuDe) validation crosslinks were violated in these models. The resultant models largely resemble the original results, although they are lower in precision (Fig. S8).

#### (5) Fit to data not used in modeling

The MHR integrative models were supported by histone deacetylation assays, mutagenesis, and co-immunoprecipitation, showing that MTA1 and the HDAC1^C^ regulate HDAC1 deacetylase activity and NuRD assembly (Pflum et al., 2001; Zhang et al., 1999). The localization of domains such as MTA1^BAH^ and RBBP4 were validated by their consistency with independently determined cryo-EM maps (Millard et al., 2020).

The MHM integrative models were supported by independent cryo-EM maps of the complex showing similar localizations for MBD2^MBD^ and MTA1^BAH^ (Millard et al., 2020). The MBD3^IDR^-MTA1^dimer^ interaction was also supported by two separate mutagenesis and co-immunoprecipitation studies (Desai et al., 2015; Millard et al., 2020).

The NuDe integrative models were corroborated by immunoprecipitation experiments showing that the MBD domain of MBD3 is buried in NuRD (Zhang et al., 1999). They were also supported by independent cryo-EM maps showing that MBD3 is proximal to MTA1^BAH^, and biochemical assays showing the importance of HDAC1^C^ interactions in NuRD (Millard et al., 2020; Pflum et al., 2001). The mapping of cancer mutations to protein-protein interfaces in the NuDe model also supported them (Fig. 6, Fig. S11, Table S1) (Forbes et al., 2006). We also validated the NuDe integrative models by an independent set of crosslinks on the endogenous NuRD complex and interacting proteins (Spruijt et al., 2021). We mapped these crosslinks onto paralogs of the subunits represented in our models. Of the 155 crosslinks in this set, 89 were within and between subunits represented in our models. All crosslinks were satisfied by the dominant cluster of the NuDe sub-complex, at a threshold of 35 Å. Finally, we also compared our model to predictions from Alphafold-Multimer (Evans et al., 2022).

##### 5.1 Mapping COSMIC mutations

We obtained a total of 356 somatic, confirmed pathogenic, point mutations for the modeled NuRD subunits (MTA1, HDAC1, RBBP4, MBD3) from the COSMIC (Catalogue of Somatic Mutations in Cancer) database (Forbes et al., 2006). For each subunit, point mutations were selected from search results based on the presence of census genes and correct documentation of current structures. To ensure the mutations studied significantly affect the function, folding, and protein-protein interaction of the protein, the “confirmed pathogenic” and “somatic” filters were applied in all cases. To test the significance of the mapping, we considered residues with 3 or more reported mutations (Alsulami et al., 2021). We generated an equal number of random mutations for each subunit and mapped them onto the NuDe integrative model. For this analysis, a mutation was considered to be at an interface if the average distance of the corresponding residue to a residue in an interacting protein is less than 5 Å.

## Data and Code availability

Files containing the input data, scripts, and output results are publicly available at https://github.com/isblab/nurd.

## Acknowledgements

We thank Vinothkumar Kutti Ragunath and lab members Satwik Pasani, Praveen Roy DS, and Varun Ullanat for useful comments on the manuscript.

Molecular graphics images were produced using the UCSF Chimera and UCSF ChimeraX packages from the Resource for Biocomputing, Visualization, and Informatics at the University of California, San Francisco (supported by NIH P41 RR001081, NIH R01-GM129325, and National Institute of Allergy and Infectious Diseases).

