## Supplementary Material for "Molecular architecture of nucleosome remodeling and deacetylase sub-complexes by integrative structure determination"

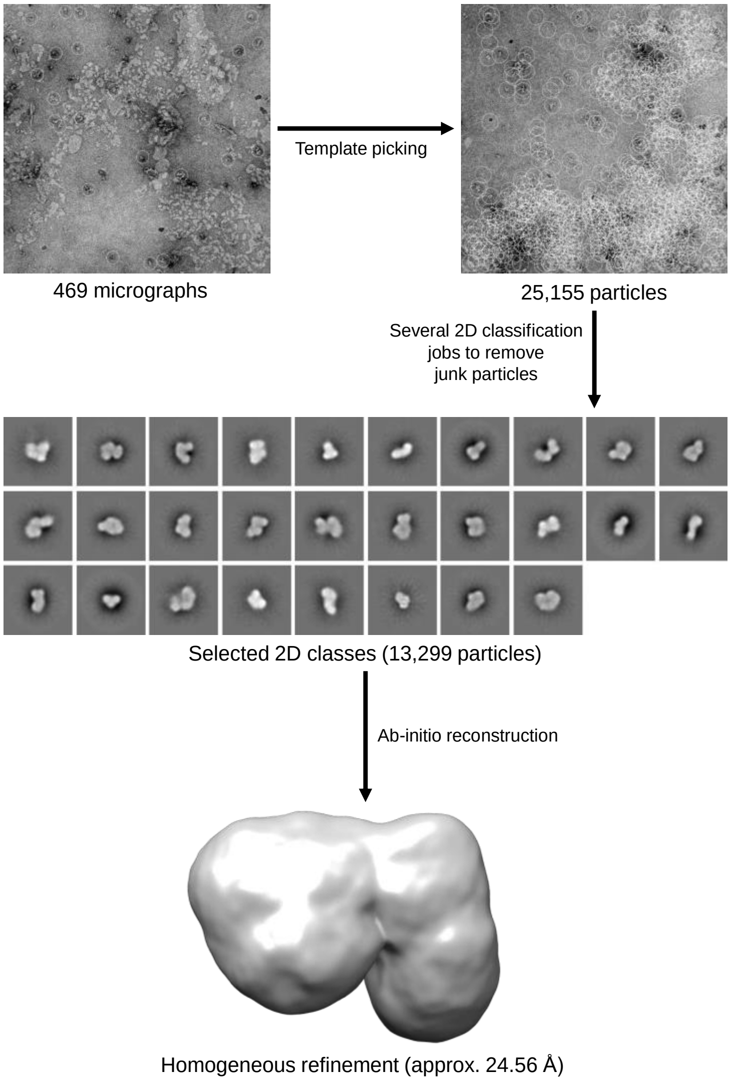


**Fig. S1 Single particle analysis of the MHR complex.** A flowthrough from the micrographs imported into Cryosparc (Punjani et al., 2017) to the final 3D reconstruction of the MHR complex using the ab-initio reconstruction job. 2D classes were generated using 50 classes per run, with particles from non-junk classes used as input to subsequent 2D classification jobs. Non-uniform refinement was not performed on the final structure due to its low resolution.

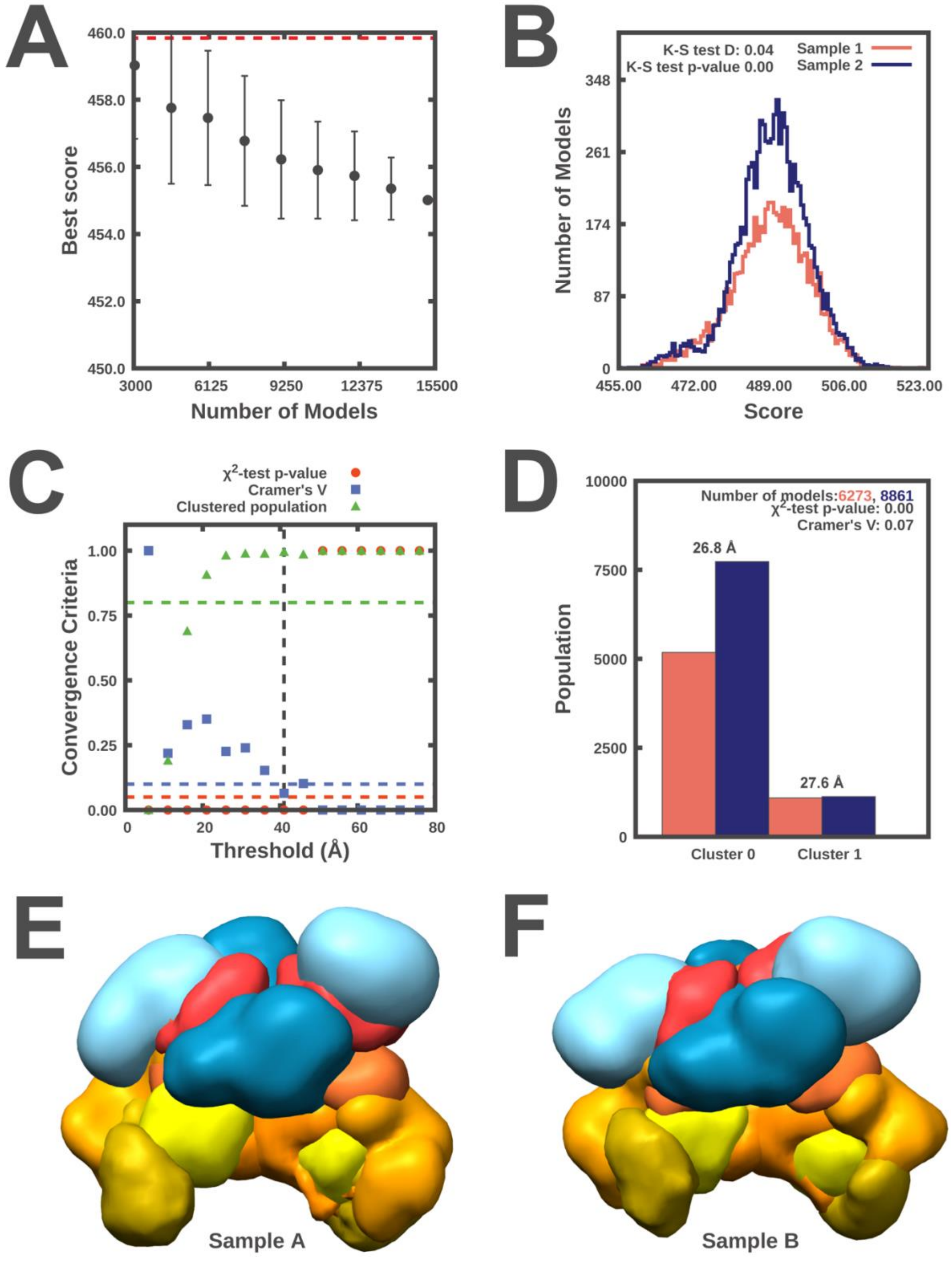


**Fig. S2 Sampling exhaustiveness protocol on MHR models** Results of test 1, convergence of the model score, for the 15200 good-scoring models; the scores do not continue to improve as more models are computed essentially independently. The error bar represents the standard deviations of the best scores, estimated by repeating sampling of models 10 times. The red dotted line indicates a lower bound reference on the total score. B. Results of test 2, testing similarity of model score distributions between samples 1 (red) and 2 (blue); the difference in the distribution of scores is significant (Kolmogorov-Smirnov two-sample test p-value less than 0.05) but the magnitude of the difference is small (the Kolmogorov-Smirnov two-sample test statistic D is 0.04); thus, the two score distributions are effectively equal. C. Results of test 3, three criteria for determining the sampling precision (Y-axis), evaluated as a function of the RMSD clustering threshold (X-axis). First, the p-value is computed using the $\chi^2$ -test for homogeneity of proportions (red dots). Second, an effect size for the $\chi^2$-test is quantified by the Cramer's V value (blue squares). Third, the population of models in sufficiently large clusters (containing at least 10 models from each sample) is shown as green triangles. The vertical dotted grey line indicates the RMSD clustering threshold at which three conditions are satisfied (p-value > 0.05 [dotted red line], Cramer's V < 0.10 [dotted blue line], and the population of clustered models > 0.80 [dotted green line]), thus defining the sampling precision of 41 Å. D. Populations of sample 1 and 2 models in the clusters obtained by threshold-based clustering using the RMSD threshold of 41 Å. Cluster precision is shown for each cluster. E. and F. Results of test 4: comparison of localization probability densities of models from sample A and sample B for the major cluster (84.95% population). The cross-correlation of the density maps of the two samples is greater than 0.96.

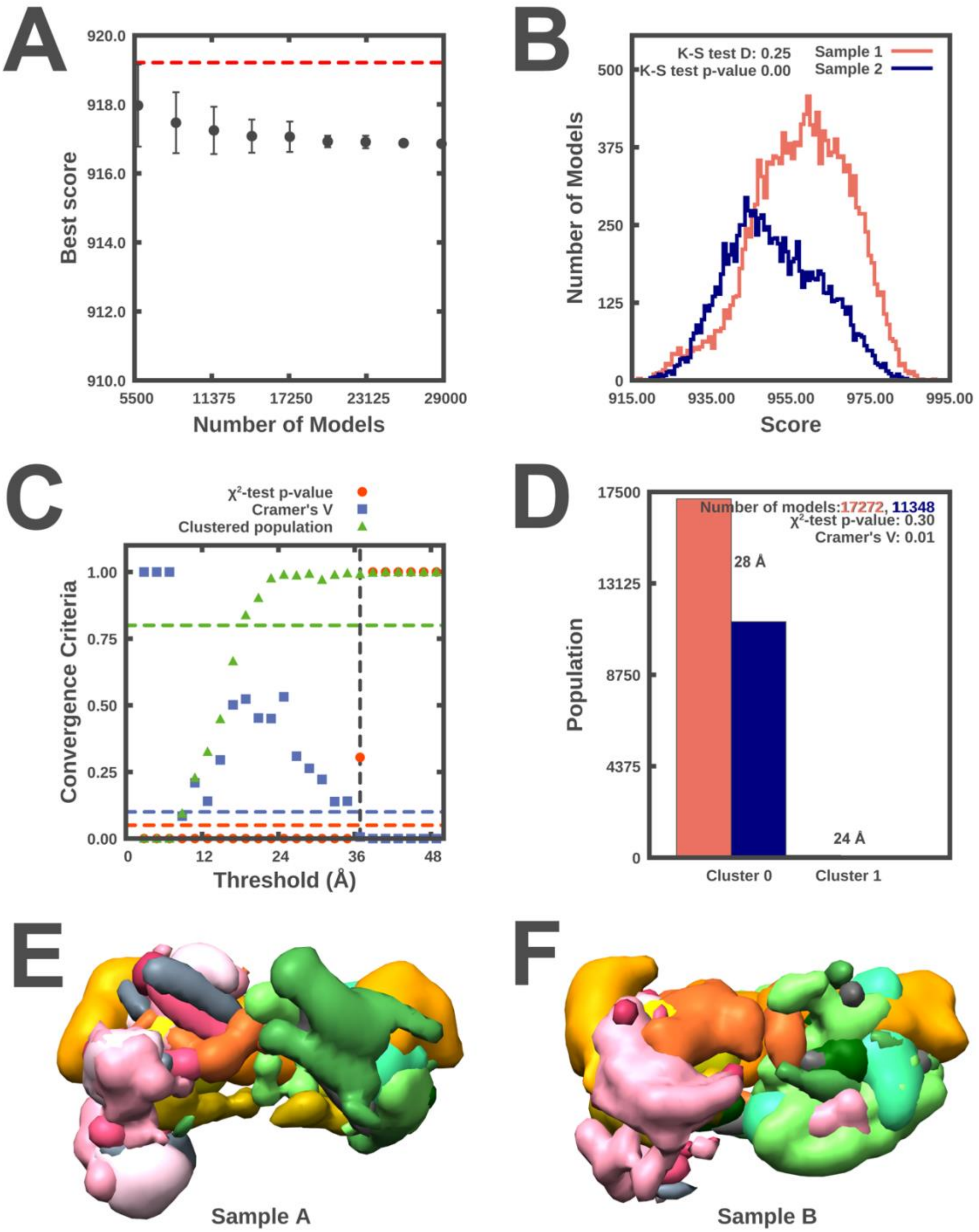


**Fig. S3 Sampling exhaustiveness protocol on MTA1$^{N}$-HDAC1-MBD3$^{GATAD2CC}$ (MHM) models** Results of test 1, convergence of the model score, for the 28836 good-scoring models; the scores do not continue to improve as more models are computed essentially independently. The error bar represents the standard deviations of the best scores, estimated by repeating sampling of models 10 times. The red dotted line indicates a lower bound reference on the total score. B. Results of test 2, testing similarity of model score distributions between samples 1 (red) and 2 (blue); the difference in the distribution of scores is significant (Kolmogorov-Smirnov two-sample test p-value less than 0.05) but the magnitude of the difference is small (the Kolmogorov-Smirnov two-sample test statistic D is 0.25); thus, the two score distributions are effectively equal. C. Results of test 3, three criteria for determining the sampling precision (Y-axis), evaluated as a function of the RMSD clustering threshold (X-axis). First, the p-value is computed using the $\chi^2$ -test for homogeneity of proportions (red dots). Second, an effect size for the $\chi^2$-test is quantified by the Cramer's V value (blue squares). Third, the population of models in sufficiently large clusters (containing at least 10 models from each sample) is shown as green triangles. The vertical dotted grey line indicates the RMSD clustering threshold at which three conditions are satisfied (p-value > 0.05 [dotted red line], Cramer's V < 0.10 [dotted blue line], and the population of clustered models > 0.80 [dotted green line]), thus defining the sampling precision of 37 Å. D. Populations of sample 1 and 2 models in the clusters obtained by threshold-based clustering using the RMSD threshold of 37 Å. Cluster precision is shown for each cluster. E. and F. Results of test 4: comparison of localization probability densities of models from sample A and sample B for the major cluster (99% population). The cross-correlation of the density maps of the two samples is 0.99.

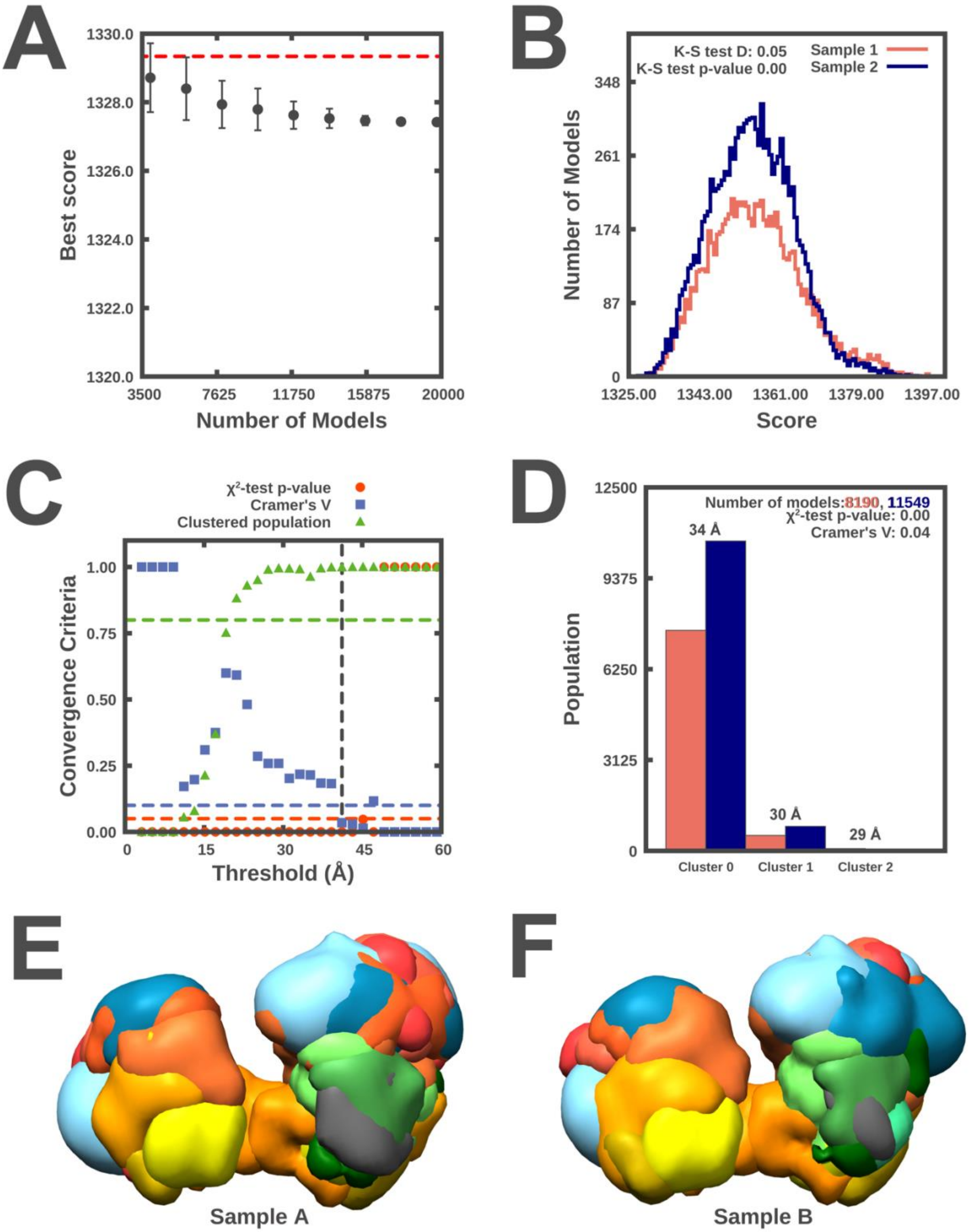


**Fig. S4 Sampling exhaustiveness protocol on NuDe integrative models** Results of test 1, convergence of the model score, for the 18005 good-scoring models; the scores do not continue to improve as more models are computed essentially independently. The error bar represents the standard deviations of the best scores, estimated by repeating sampling of models 10 times. The red dotted line indicates a lower bound reference on the total score. B. Results of test 2, testing similarity of model score distributions between samples 1 (red) and 2 (blue); the difference in the distribution of scores is significant (Kolmogorov-Smirnov two-sample test p-value less than 0.05) but the magnitude of the difference is small (the Kolmogorov-Smirnov two-sample test statistic D is 0.05); thus, the two score distributions are effectively equal. C. Results of test 3, three criteria for determining the sampling precision (Y-axis), evaluated as a function of the RMSD clustering threshold (X-axis). First, the p-value is computed using the $\chi^2$ -test for homogeneity of proportions (red dots). Second, an effect size for the $\chi^2$-test is quantified by the Cramer's V value (blue squares). Third, the population of models in sufficiently large clusters (containing at least 10 models from each sample) is shown as green triangles. The vertical dotted grey line indicates the RMSD clustering threshold at which three conditions are satisfied (p-value > 0.05 [dotted red line], Cramer's V < 0.10 [dotted blue line], and the population of clustered models > 0.80 [dotted green line]), thus defining the sampling precision of 41 Å. D. Populations of sample 1 and 2 models in the clusters obtained by threshold-based clustering using the RMSD threshold of 41 Å. Cluster precision is shown for each cluster. E. and F. Results of test 4: comparison of localization probability densities of models from sample A and sample B for the major cluster (71% population). The cross-correlation of the density maps of the two samples is 0.99.

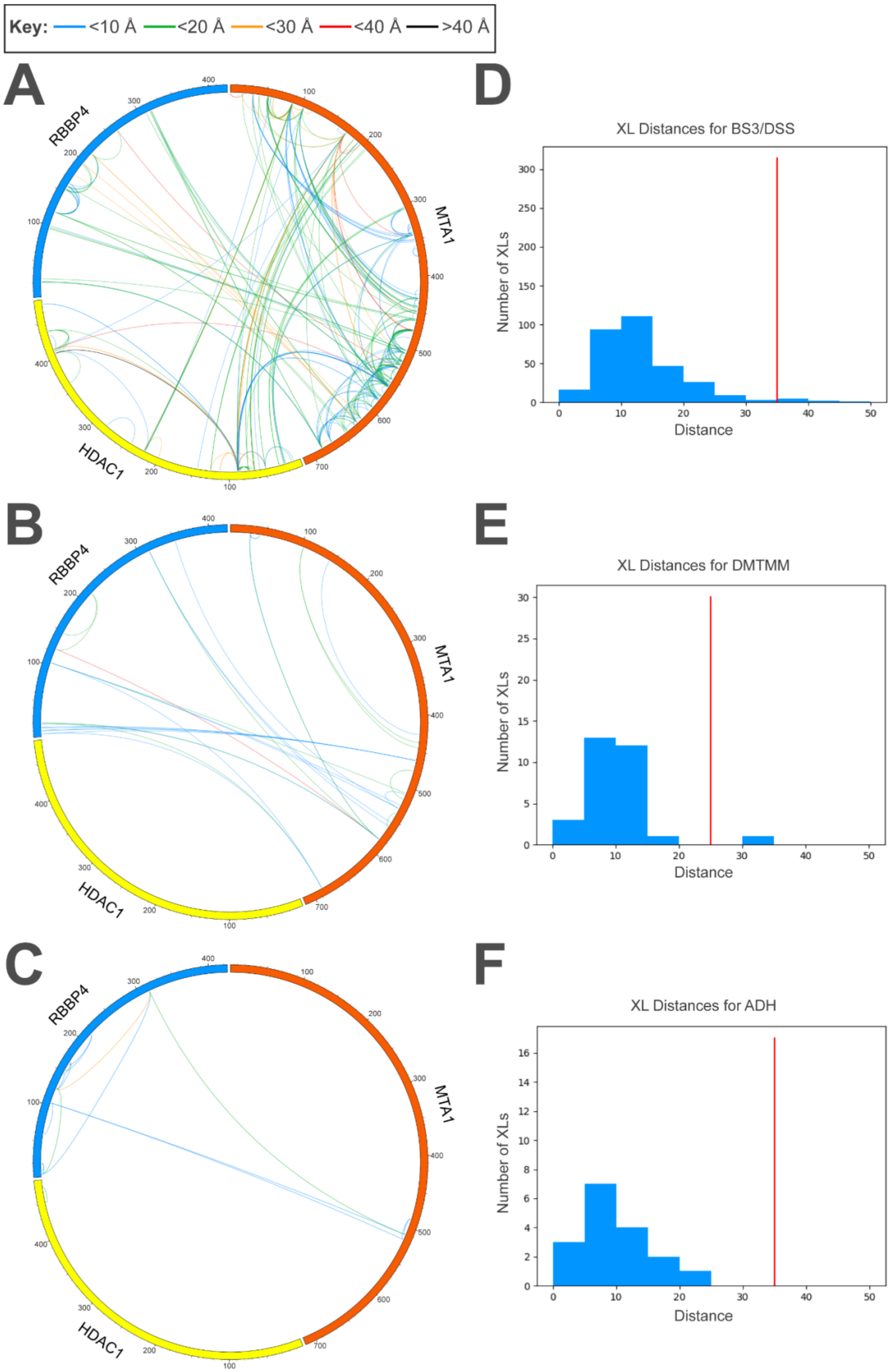

Key: <10 Å <20 Å <30 Å <40 Å >40 Å
A
RBBP4
MTA1
HDAC1
B
RBBP4
MTA1
HDAC1
C
RBBP4
MTA1
HDAC1
D
XL Distances for BS3/DSS
Number of XLs
Distance
E
XL Distances for DMTMM
Number of XLs
Distance
F
XL Distances for ADH
Number of XLs
Distance

**Fig. S5 Results of crosslink fitting for MHR models** CX-CIRCOS (http://cx-circos.net/) plots are shown for A. BS3/DSS, B. DMTMM, and C. ADH crosslinks on the ensemble of MHR models from the major cluster. Each line depicts a crosslink; its color depicts the minimum distance between the corresponding crosslinked residues in the ensemble, as shown in the color key (top). Histograms showing the distribution of the minimum crosslink distance in the ensemble for D. BS3/DSS, E. DMTMM, and F. ADH crosslinks. The red line indicates the distance threshold for a crosslink type.

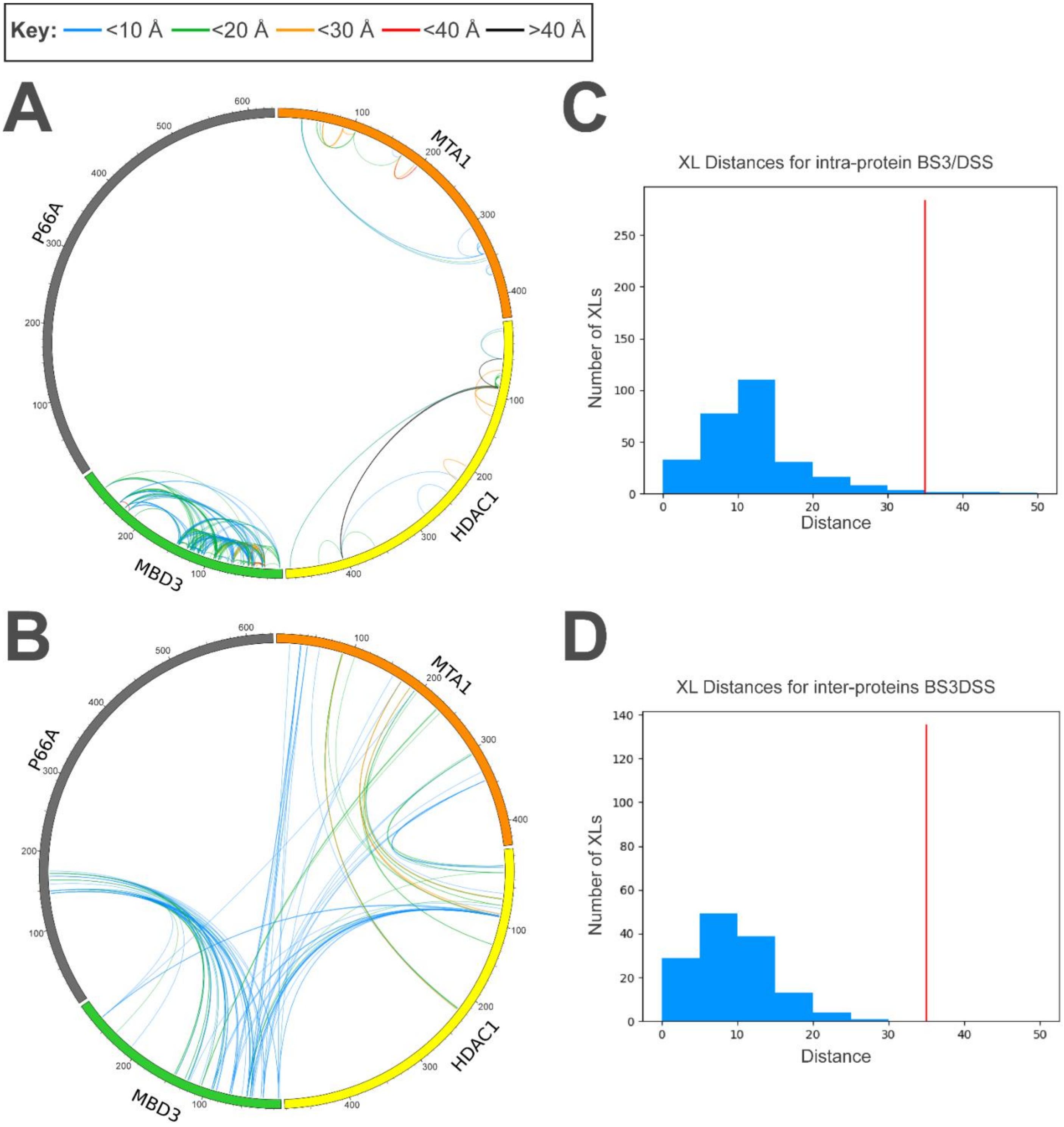


**Fig. S6 Results of crosslink fitting for MTA1$^{N}$-HDAC1-MBD3$^{GATAD2CC}$ (MHM) models** CX-CIRCOS (http://cx-circos.net/) plots are shown for A. BS3/DSS, B. DMTMM, and C. ADH crosslinks on the ensemble of MHM models from the major cluster. Each line depicts a crosslink; its color depicts the minimum distance between the corresponding crosslinked residues in the ensemble, as shown in the color key (top). Histograms showing the distribution of the minimum crosslink distance in the ensemble for D. BS3/DSS, E. DMTMM, and F. ADH crosslinks. The red line indicates the distance threshold for a crosslink type.

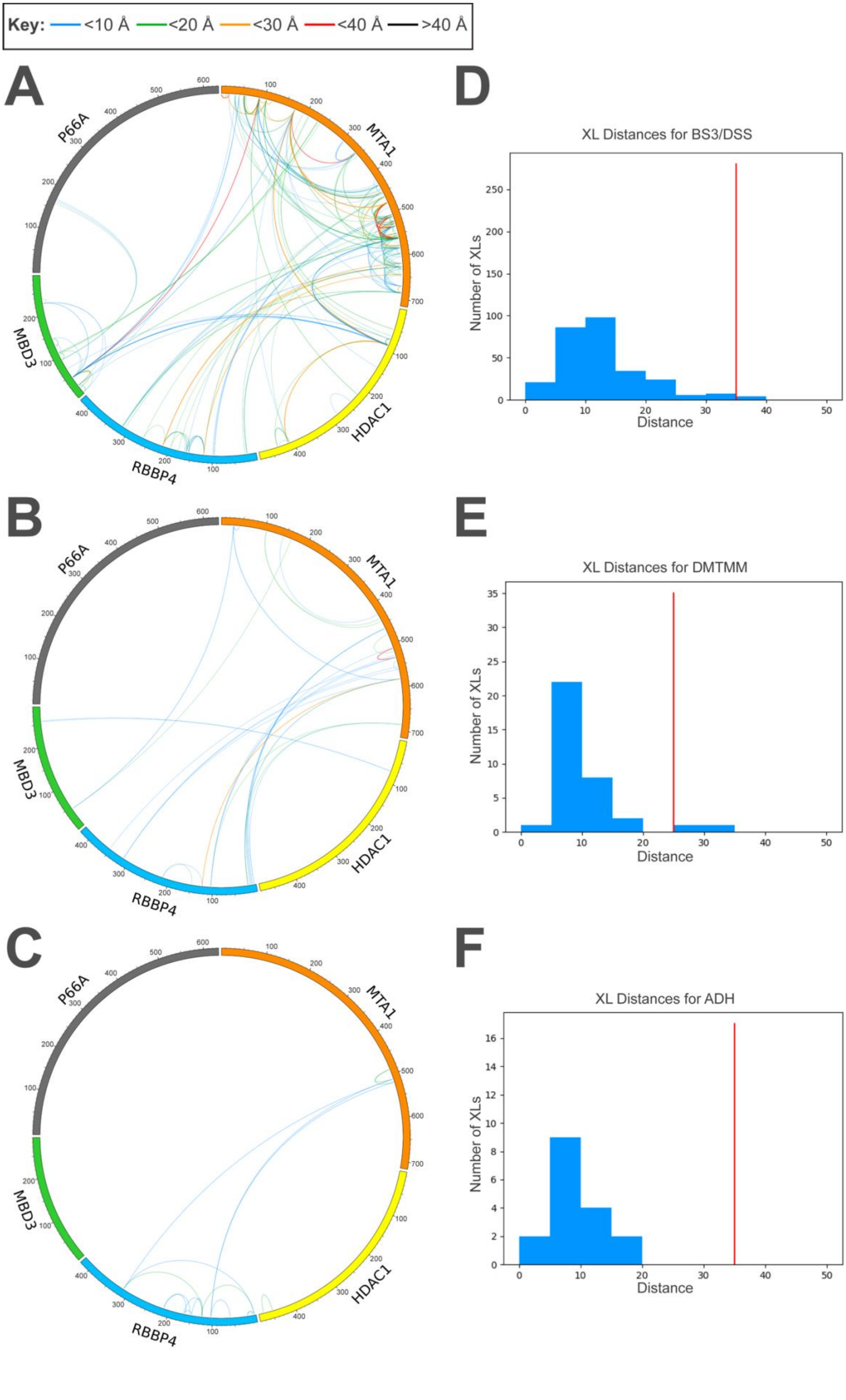
Key: <10 Å <20 Å <30 Å <40 Å >40 Å
A
B
C
P66A
MTA1
MBD3
RBBP4
HDAC1
D
XL Distances for BS3/DSS
E
XL Distances for DMTMM
F
XL Distances for ADH
Number of XLs
Distance

**Fig. S7 Results of crosslink fitting for NuDe models** CX-CIRCOS (http://cx-circos.net/) plots are shown for A. BS3/DSS, B. DMTMM, and C. ADH crosslinks on the ensemble of NuDe models from the major cluster. Each line depicts a crosslink; its color depicts the minimum distance between the corresponding crosslinked residues in the ensemble, as shown in the color key (top). Histograms showing the distribution of the minimum crosslink distance in the ensemble for D. BS3/DSS, E. DMTMM, and F. ADH crosslinks. The red line indicates the distance threshold for a crosslink type.

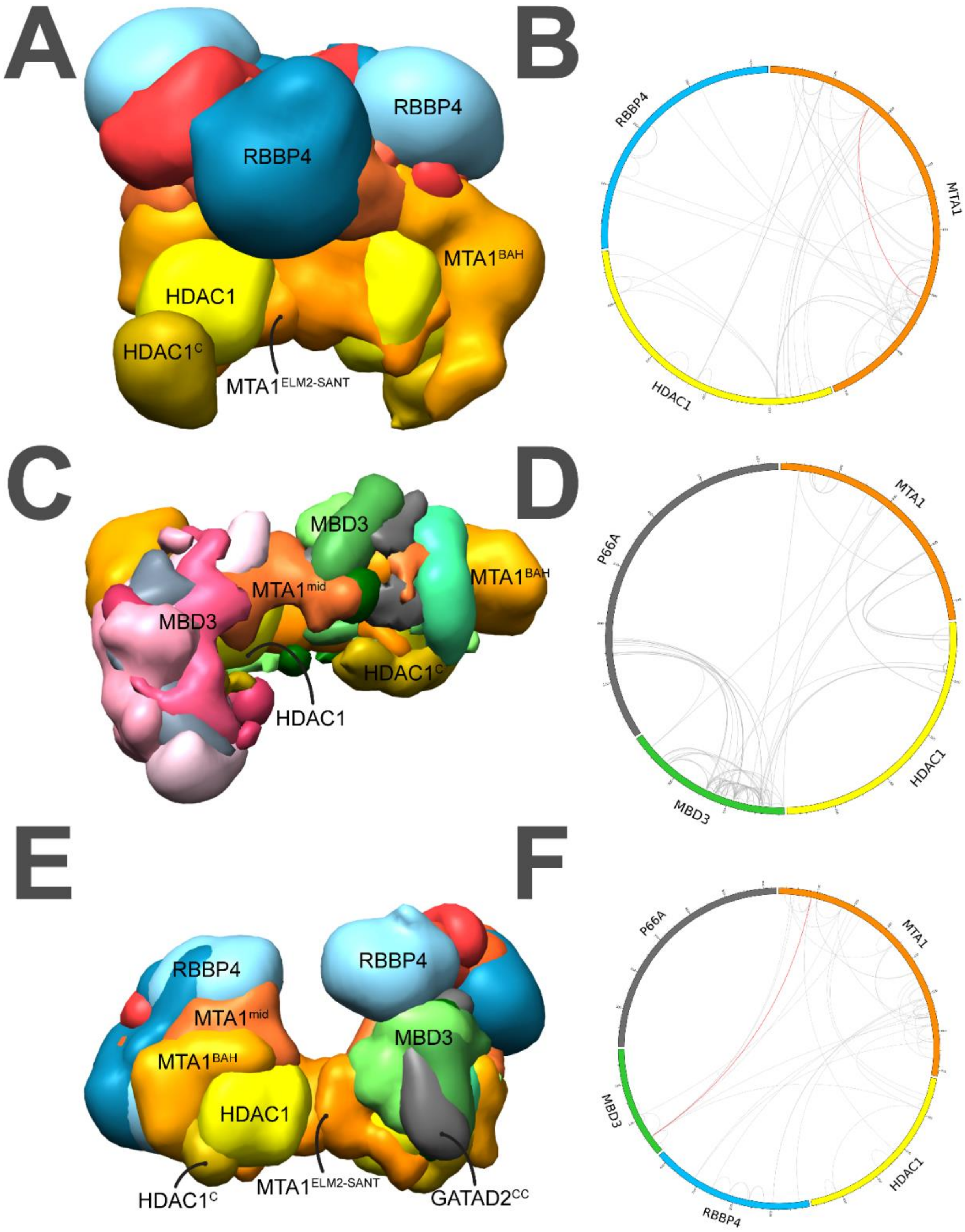

**Fig. S8 Results of jack-knifing tests** MHR, MHM, and NuDe models were recomputed by omitting a random subset of the input BS3/DSS crosslinks. The recomputed models were validated by their fit to the omitted crosslinks. Localization probability densities for the ensemble of recomputed models from the top cluster for A. MHR, C. MHM, and E. NuDe sub-complexes. Map colors and contour levels are similar to the main runs (Fig. 3- Fig. 5). CX-CIRCOS (http://cx-circos.net/) plots showing fit to the omitted crosslinks for the recomputed B. MHR, D. MHM, and F. NuDe models. Grey (red) lines indicate satisfied (violated) crosslinks.

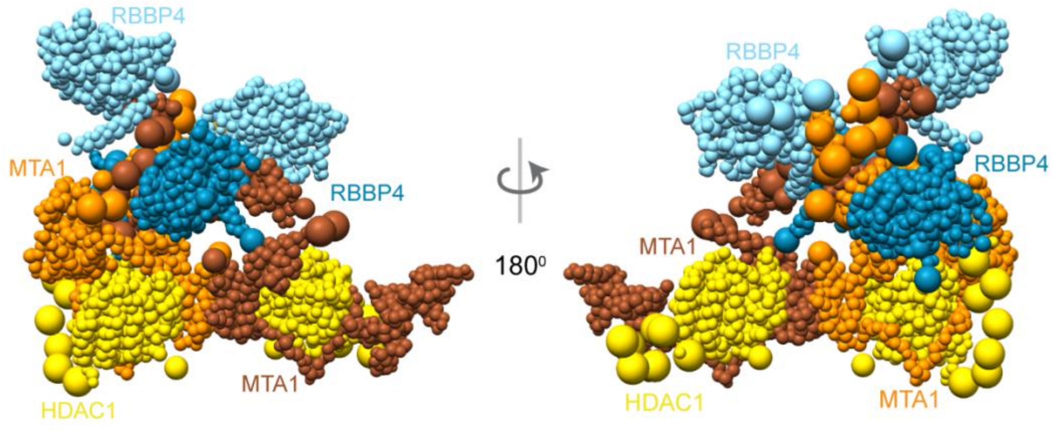


**Fig. S9 Integrative model of the MHR complex based on crosslinks alone** Representative bead model from the most populated cluster of analyzed integrative models for the MHR complex, colored by subunit. The two copies of MTA1 are shown in different colors (orange and brown) to illustrate the crossover.

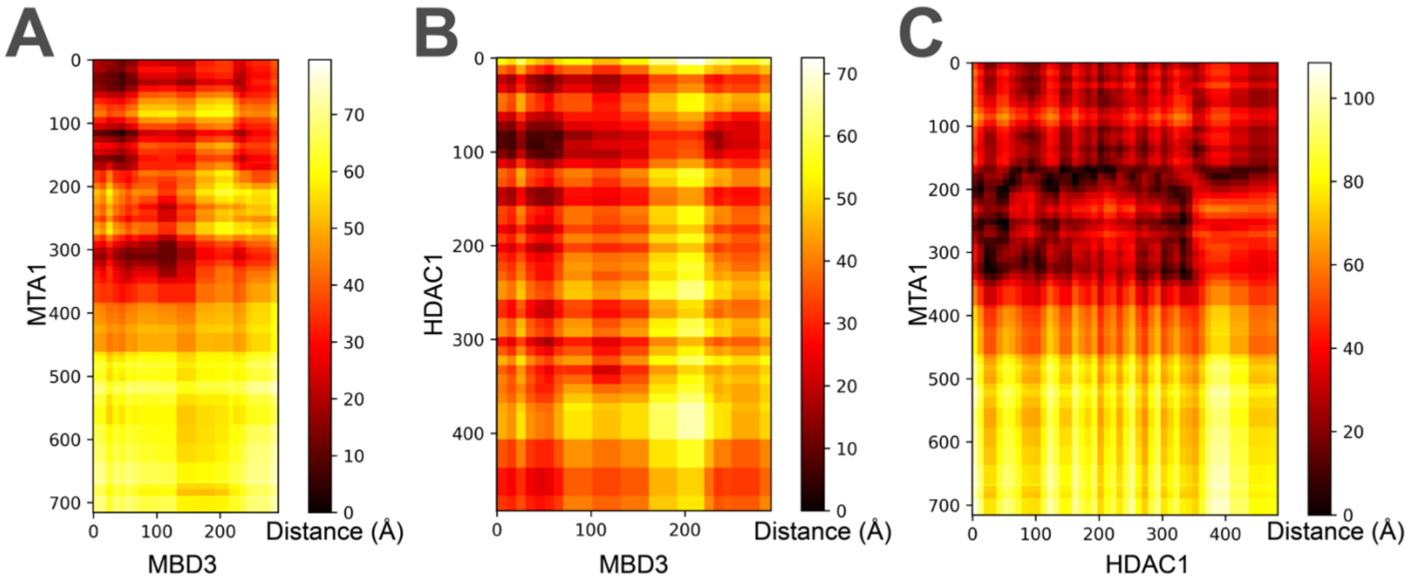


**Fig. S10 Distance maps for protein interactions in NuDe**. The distance maps show the average pairwise residue distances in the ensemble of NuDe models for the A. MBD3-MTA1, B. MBD3-HDAC1, and C. HDAC1-MTA1 protein pairs. For a pair of residues, the map indicates the distance between the surfaces of the corresponding beads averaged over the ensemble.

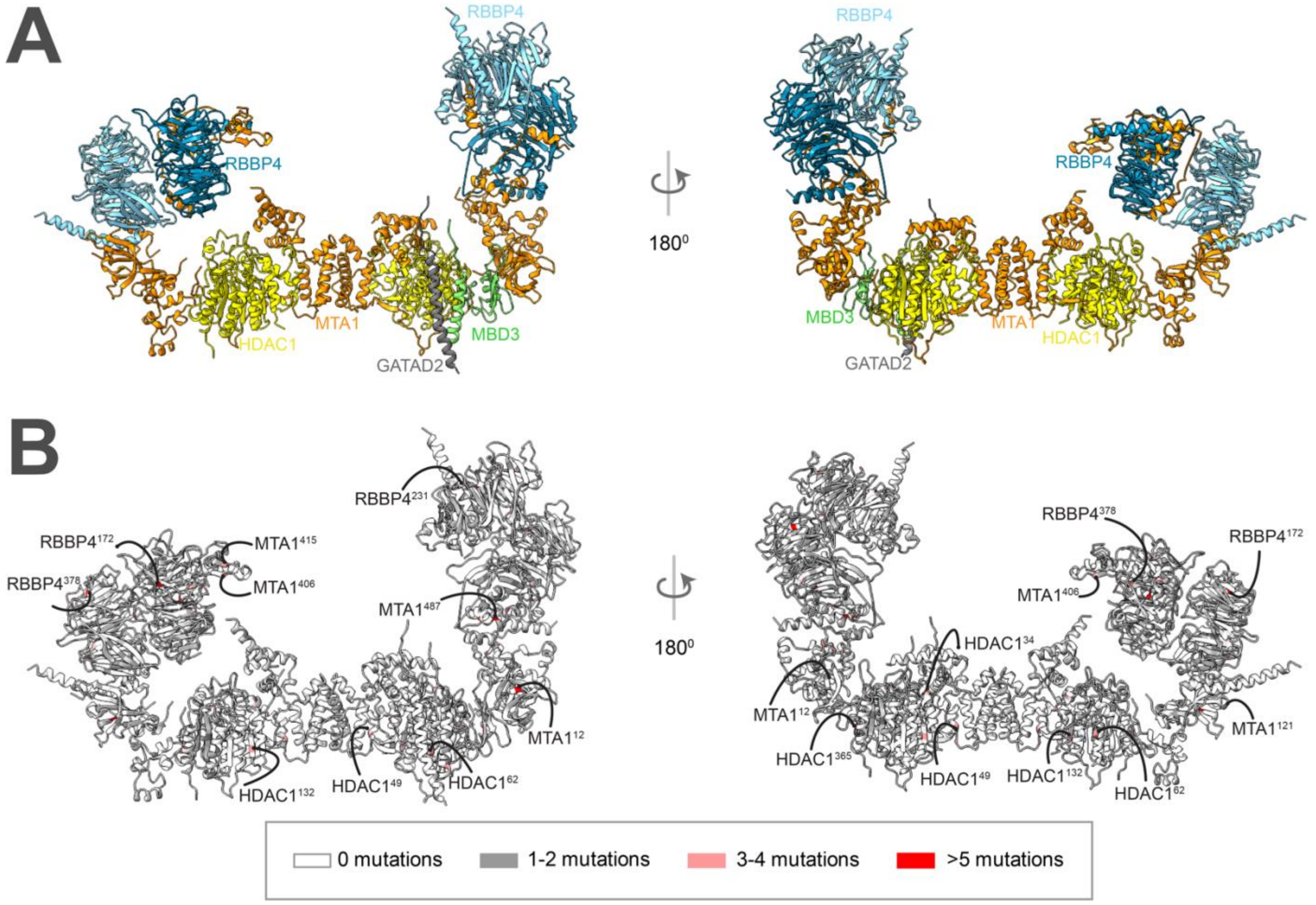


**Fig. S11 COSMIC mutations on the structured regions of NuDe complex** Somatic, confirmed pathogenic, point mutations from the COSMIC database (Forbes et al., 2006) mapped onto regions with known structure in the NuDe integrative model shown in Fig. 4A. A. Regions with known atomic structure in the integrative model colored by subunit. B. Mutations on residues in regions of known structure, colored according to the legend.

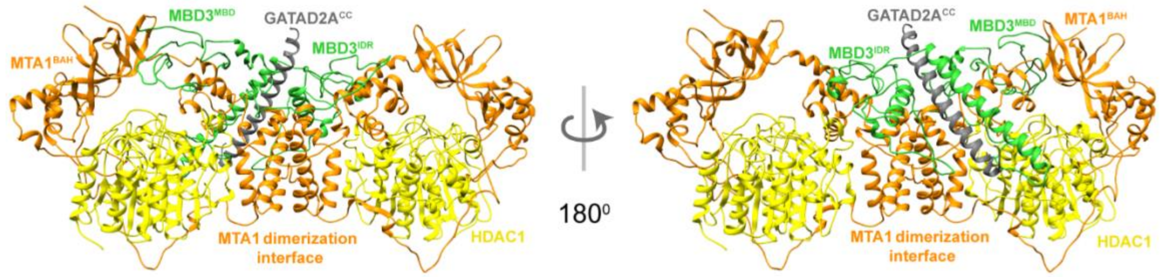


**Fig. S12 AlphaFold multimer (Evans et al., 2022) predictions for 2:2:1:1 MTA1:HDAC1:MBD3:GATAD2A$^{CC}$ complex** Front and back views of the top ranked model predicted by AlphaFold 2 multimer showing the proximity of MBM3$^{MBD}$ to MTA1$^{BAH}$ and the MBD3$^{IDR}$-MTA1 dimerization interface.

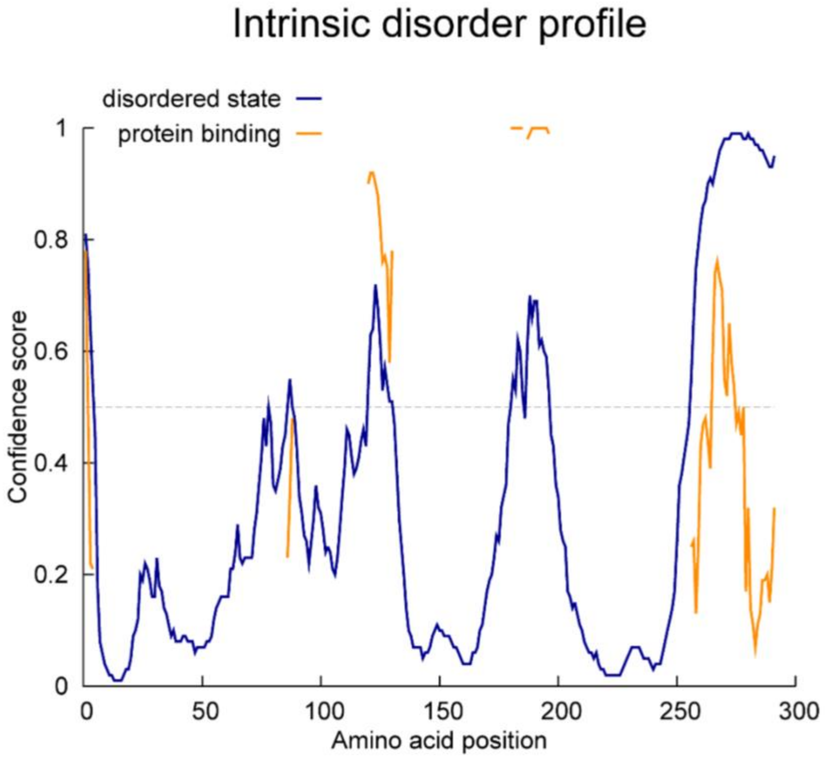


**Fig. S13 Disorder prediction for MBD3** PONDR® (http://www.pondr.com) (Romero et al., 1997, 2001) disorder prediction for MBD3.

**Table S1. COSMIC mutations on the NuDe integrative model** Somatic, confirmed pathogenic point mutations from the COSMIC database [(Forbes et al., 2006)] mapped on the NuDe integrative model. A. Mutations in previously undescribed protein-protein interfaces in the model. Residues from two proteins are at an interface if the average distance between their corresponding bead surfaces is less than 5 Å in the cluster of models. B. Mutations in exposed binding sites between modeled proteins and known binding partners, based on the representative NuDe model.

| **A. COSMIC mutations in previously undescribed protein-protein interfaces in NuDe** | | | |
|---|---|---|---|
| **NuRD protein domain** | **Residues with mutations** | **Possible interacting partners in NuRD** | **Number of COSMIC mutations** |
| MBD3$^{MBD}$ | 2, 7, 12, 14, 19, 21, 29, 31, 39, 41, 43, 45, 55, 65, 68 | MTA1$^{BAH}$, HDAC1$^{60-100}$ | 1-2 |
| | 17, 60 | MTA1$^{BAH}$, HDAC1$^{60-100}$ | 3-4 |
| MTA1$^{BAH}$ | 10, 15, 24, 31, 46, 53, 60, 63, 81, 88, 90, 104, 107, 112, 121, 128, 133, 135, 143, 146, 153, 155, 158 | MBD3$^{MBD}$, HDAC1$^{C}$ | 1-2 |
| | 12 | MBD3$^{MBD}$, HDAC1$^{C}$ | 5+ |
| | 61, 64, 69, 73, 80, 86, 94, 96 | MBD3$^{MBD}$ | 1-2 |
| HDAC1$^{60-100}$ | 62 | MBD3$^{MBD}$ | 3-4 |
| HDAC1$^{C}$ | 376, 379, 388, 391,396, 404, 411, 420, 423, 425, 431, 435, 447, 462, 474, 476 | MTA1$^{BAH}$ | 1-2 |
| | 398 | MTA1$^{BAH}$ | 3-4 |

| **B. COSMIC mutations in exposed binding sites to known interactors in NuDe** | | | |
|---|---|---|---|
| **NuRD protein** | **Residues with mutations** | **Associated partner outside NuRD if known** | **Number of COSMIC mutations** |
| RBBP4 | 39, 46, 71, 75, 129, 321,376, 396, 398 | H3, FOG1/2, other ZF containing TFs [(Lejon et al., 2011; Liu et al., 2015; Moody et al., 2018; Schmidberger et al., 2016)] | 1-2 |
| RBBP4 | 40,378 | H3, FOG1/2, other ZF containing TFs [(Lejon et al., 2011; Liu et al., 2015; Moody et al., 2018; Schmidberger et al., 2016)] | 3-4 |
| MTA1$^{BAH}$ | 46, 53, 153, 155, 158 | Nucleosome (based on Sir3-BAH **PDB** 3TU4 [6]), MAT1 [(Millard et al., 2014)] | 1-2 |
| MTA1$^{mid}$ | 390, 394, 397, 401, 410, 416, 420, 424 | MAT1 [(Mazumdar et al., 2001)] | 1-2 |
| MTA1$^{mid}$ | 406, 415 | MAT1 [(Mazumdar et al., 2001)] | 3-4 |
| MTA1$^{mid}$ | 372 | NRIF3 [(Talukder et al., 2004)] | 5+ |
| MTA1$^{USR}$ | 547, 550, 562, 564, 578, 581, 601, 603, 606, 612, 617, 619, 633, 647, 652, 658, 666 | MICoA [(Mishra et al., 2003)] | 1-2 |
| MTA1$^{USR}$ | 549, 566, 571 | MICoA [(Mishra et al., 2003)] | 3-4 |
| MTA1$^{C}$ | 697, 699, 705, 709, 715 | MICoA [(Mishra et al., 2003)] | 1-2 |
| HDAC1 active site | 140, 150, 303 | Histone tails, TFs [(Scafuri et al., 2020)] | 1-2 |
| MBD3$^{IDR}$ | 77, 78, 80, 85, 87, 98, 103, 107 | Evi1 [(Spensberger & Delwel, 2008)], Aurora A [(Sakai et al., 2002)] | 1-2 |
